# Dissecting the molecular basis for the modulation of neurotransmitter GPCR signaling by GINIP

**DOI:** 10.1101/2023.04.20.537566

**Authors:** Alex Luebbers, Myles Zhou, Stephen J Eyles, Mikel Garcia-Marcos

**Author notes:** Corresponding author: Mikel Garcia-Marcos.

## Abstract

It is well-established that activation of heterotrimeric G-proteins (Gαβγ) by G-protein-coupled receptors (GPCRs) stimulated by neurotransmitters is a key mechanism underlying neuromodulation. Much less is known about how G-protein regulation after receptor-mediated activation contributes to neuromodulation. Recent evidence indicates that the neuronal protein GINIP shapes GPCR inhibitory neuromodulation via a unique mechanism of G-protein regulation that controls neurological processes like pain and seizure susceptibility. However, the molecular basis of this mechanism remains ill-defined because the structural determinants of GINIP responsible for binding Gαi subunits and regulating G-protein signaling are not known. Here, we combined hydrogen-deuterium exchange mass-spectrometry, protein folding predictions, bioluminescence resonance energy transfer assays, and biochemical experiments to identify the first loop of the PHD domain of GINIP as an obligatory requirement for Gαi binding. Surprisingly, our results support a model in which GINIP undergoes a long-range conformational change to accommodate Gαi binding to this loop. Using cell-based assays, we demonstrate that specific amino acids in the first loop of the PHD domain are essential for the regulation of Gαi-GTP and free Gβγ signaling upon neurotransmitter GPCR stimulation. In summary, these findings shed light onto the molecular basis for a post-receptor mechanism of G-protein regulation that fine-tunes inhibitory neuromodulation.

## INTRODUCTION

Metabotropic neurotransmitter responses in the nervous system are largely mediated by G protein-coupled receptors (GPCRs) (Roth, 2019). GPCRs are the largest family of membrane receptors in the human genome and are capable of relaying signals from a wide range of stimuli, including the vast majority of neurotransmitters (Weis and Kobilka, 2018). In contrast to neurotransmitter ionotropic receptors that rapidly and directly control ion fluxes in neurons, the control of electrical responses by GPCRs occurs at a slower timescale due to the involvement of intermediary steps through heterotrimeric G proteins and second messengers (Greengard, 2001; Roth, 2019; Zurawski et al., 2019). Thus, a critical role of GPCRs in the nervous system is to exert neuromodulatory effects over rapid ionotropic responses. This neuromodulatory function is not only critical for the proper processing of complex neurochemical signals, but also makes GPCRs an attractive target to develop drugs that correct neurotransmission imbalances associated with neurological and neuropsychiatric disorders (Hauser et al., 2017; Hopkins and Groom, 2002; Santos et al., 2017; Sriram and Insel, 2018). By having more nuanced effects in neurotransmission, GPCR-targeting agents hold the promise of achieving efficacy with reduced undesired effects. This promise is well supported by the fact that GPCRs are the target for over one-third of clinically used drugs (Hauser et al., 2017), including many medications for neurological and neuropsychiatric disorders like schizophrenia, bipolar disorder, depression, or Parkinson’s disease, among others.

Understanding the molecular basis of GPCR-mediated signaling is critical to understand the mechanism of action for both endogenous neurotransmitters as well as pharmacological agents. GPCRs activate heterotrimeric G proteins (Gαβγ), which are divided in four families (G_s_, G_i/o_, G_q/11_, G_12/13_) based on sequence similarity and function (Gilman, 1987). Among them, G_i/o_ family proteins mediate neuroinhibitory signals, such as those triggered by many neurotransmitter receptors, like the GABA_B_ receptor (GABA_B_R), dopamine 2 receptor (D2R), or α2 adrenergic receptors (α2-AR), among others (Bylund et al., 1994; Gurevich et al., 2016; Ulrich and Bettler, 2007). These receptors promote the exchange of GDP for GTP on the Gαi-subunits, resulting in adoption of an active conformation than binds to adenylyl cyclase and inhibit its ability to produce the second messenger cAMP. In parallel, this also triggers the dissociation (or rearrangement) of Gβγ dimers, which in turn modulate a number of different downstream effectors, including ion channels and components of the exocytic machinery (Betke et al., 2012; Csanady, 2017). Signaling is turned off upon hydrolysis of GTP, which leads to the reassociation of Gαi and Gβγ into an inactive heterotrimer. This core G protein cycle mechanisms is influenced by a growing number of cytoplasmic proteins that modulate nucleotide handling by the Gα subunit, thereby having a profound effect on the lifetime or amplitude of G protein signaling. For example, Regulators of G protein Signaling (RGS) proteins are <u>G</u>TPase <u>A</u>ctivating <u>P</u>roteins (GAPs) that accelerate nucleotide hydrolysis, resulting in cessation of signaling and reformation of the inactive heterotrimer (Berman et al., 1996; De Vries et al., 1996; Dohlman and Thorner, 1997; Soundararajan et al., 2008; Watson et al., 1996). In the nervous system, RGS proteins have profound effects in neurotransmitter signaling and neurological behaviors (Anderson et al., 2010; Han et al., 2006; Neubig and Siderovski, 2002; Zhou et al., 2012), highlighting the importance of post-receptor regulatory mechanisms.

Recent work has established a unique mechanism by which the neuronal protein GINIP acts as a broad regulator of G_i_-coupled GPCR signaling in neurons that alters the balance of signaling after receptor activation (Park, 2023). GINIP binds to Gαi-GTP, but does not affect directly how nucleotides are handled by the G protein. Instead, GINIP prevents the engagement of Gαi with adenylyl cyclase and the subsequent modulation of cAMP levels. Simultaneously, GINIP prevents binding of Gαi-GTP to RGS GAPs, which extends the lifetime of free Gβγ signaling. This mechanism is essential for preventing imbalances of neurotransmission, as GINIP knock-out mice display higher susceptibility to seizures and increased pain in different experimental paradigms (Gaillard et al., 2014). Despite these critical roles of GINIP in GPCR-mediated neuromodulation, the molecular basis for it remains ill-defined because the structural determinants of GINIP responsible for binding GTP-bound Gαi and regulating G-protein signaling are not known. Here, we define a structural determinant of GINIP that is essential for binding to active Gαi and for regulating both Gαi-GTP and free Gβγ signaling in response to GPCR stimulation. Our results strongly suggest that the mechanism by which GINIP regulates neurotransmitter signaling involves an effector-like binding mode on the α3/SwII groove of active Gαi, which helps explain its functional antagonism of adenylyl cyclase and RGS protein engagement with G proteins.

## RESULTS

### GINIP forms a stable one-to-one complex with Gαi3-GTP

Previous work has established that GINIP binds with high affinity (i.e., K_D_ ∼65 nM) to active Gαi3 (Park, 2023), but the specific regions of GINIP involved in this protein-protein interaction were not elucidated. To identify these regions, we set out to carry out hydrogen-deuterium exchange mass spectrometry (HDX-MS) experiments with GINIP in the presence and absence of Gαi3. We reasoned that Gαi3 binding would alter the dynamics of GINIP regions directly involved in the physical interaction, as well as other regions affected indirectly by binding, which would manifest as changes in the exchange of protons with the solvent detected by HDX-MS (Engen and Wales, 2015; Englander, 2006; Hvidt and Nielsen, 1966; James et al., 2022). When purified Gαi3 activated by loading with the non-hydrolyzable GTP analog GTPγS was incubated with purified GINIP at approximately equimolar protein amounts, a one-to-one GINIP:Gαi3 complex could be recovered after size-exclusion chromatography (**Fig. 1A**), suggesting that the stability of this complex might be suitable for HDX-MS. Next, we validated the stability of the GINIP:Gαi3 complex under conditions identical to those planned for HDX-MS experiments. First, we formed the complex by mixing GINIP and Gαi3 at a 1:2 ratio to ensure that all GINIP is Gαi3-bound, a critical consideration for HDX-MS (Masson et al., 2019), and analyzed its stability by gel-filtration chromatography during the span of 24 hours after formation, which would cover the duration of the HDX-MS experiments. We found that the GINIP:Gαi3 complex remained intact after 24 h (**Fig. 1B**), validating appropriate conditions for the HDX-MS experiments. While GINIP has been described to be myristoylated and to possess putative binding sites for Zn^2+^ and Ca^2+^, these experiments were carried out with non-myristoylated GINIP without cation supplementation because our results indicated that neither myristoylation nor divalent cations significantly affected Gαi3 binding by GINIP (**Fig. S1**).

**Figure 1.**
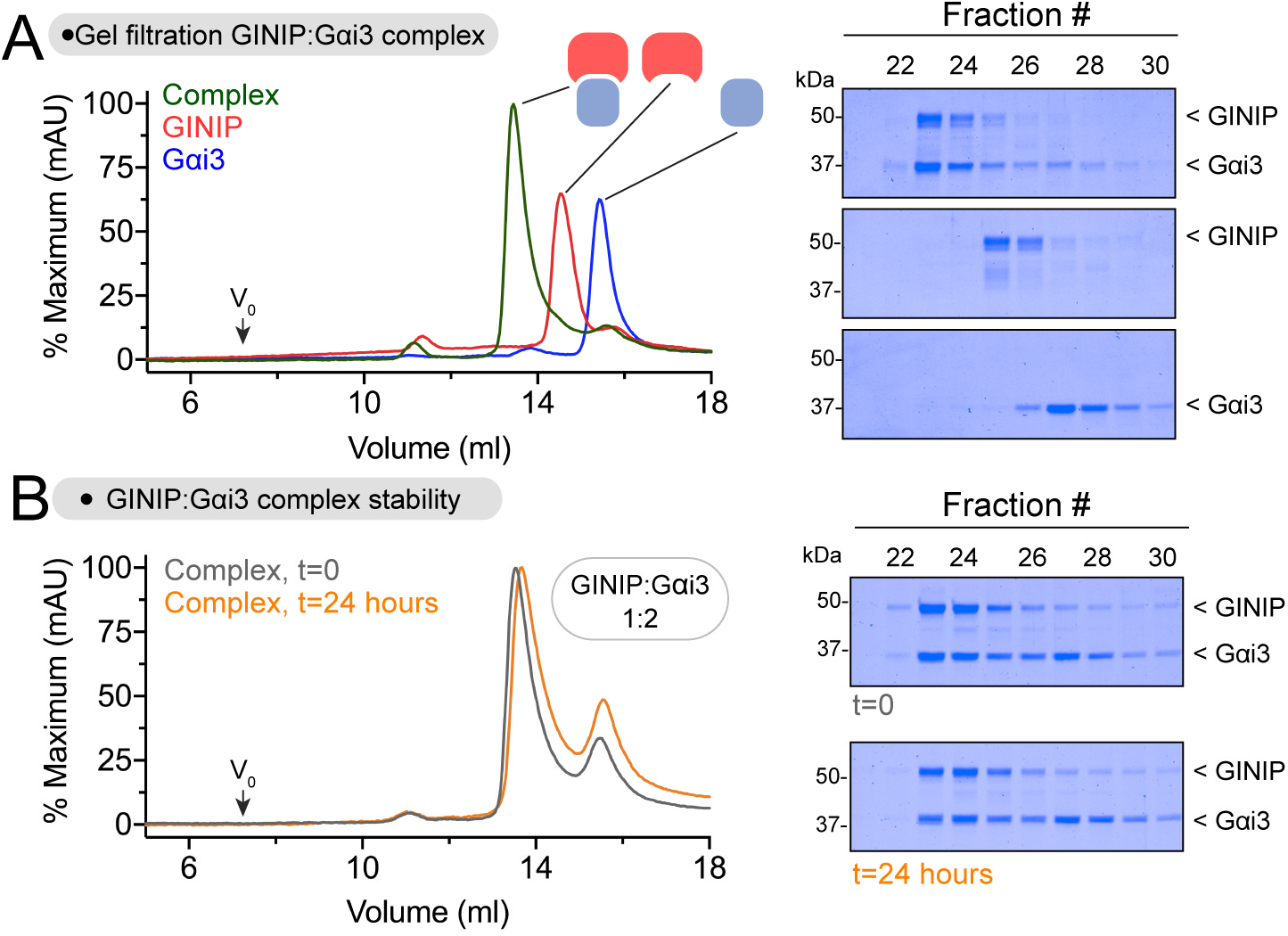
GINIP forms a stable equimolar complex with Gαi3. **(A)** Elution profiles of GINIP and Gαi3 alone or after forming a complex. *Left,* overlay of gel filtration chromatography curves for GINIP:Gαi3-GTPγS (green), GINIP (red), and Gαi3-GTPγS (blue) run on an Superdex S200 column. *Right,* Coomassie-stained gels showing selected fractions from the gel filtration chromatography. **(B)** The GINIP:Gαi3 complex is stable for >24 hours at 4 °C. *Left,* overlay of gel filtration chromatography curves for a GINIP:Gαi3-GTPγS complex formed by mixed the individual species in a 1:2 molar ratio for 15 minutes (t=0, grey) or for 24 hours (t=24 hours, orange) before application to a Superdex S200 column. *Right,* Coomassie-stained gels showing selected fractions from the gel filtration chromatography. Representative gel images from 2 independent experiments are shown.

### Gαi3 binding alters the dynamics of discrete protein regions in GINIP

GINIP alone or 1:2 GINIP-Gαi3 complexes prepared as described in the previous section were processed in a system specifically outfitted for HDX-MS. Briefly, samples were diluted in deuterated buffer for different times (10, 100, 1000, 10000 s), followed by rapid quenching and in-line digestion by pepsin before chromatographic separation and MS detection (**Fig. 2A**). Relative deuterium uptake was determined by mass analysis of peptide spectra across time points (Masson et al., 2019). Differences in deuterium uptake in GINIP caused by Gαi3 were determined by subtractive analysis of uptake data for the GINIP-Gαi3 complex relative to GINIP alone (**Fig. 2B**, **Fig. S2**). This subtractive analysis revealed a few regions with a marked decrease in deuterium uptake (approx. – 20% uptake) but also one region with a marked increase (approx. + 20% uptake). Uptake plots for representative peptides of these regions are shown in **Fig. 2C**. Because ligand binding typically results in a decrease of deuterium uptake in the region of direct physical engagement due to reduced local dynamics and exchange with the solvent, we initially focused our attention on these type of regions to identify Gαi3 binding sites on GINIP. However, protein-protein binding experiments revealed that mutations in amino acids of four different regions of GINIP displaying reduced deuterium uptake did not have a significant effect on binding active Gαi3 (**Fig. S3**). Taken together, these results suggest that, even though Gαi3 causes clear changes in the dynamics of discrete regions of GINIP, the binding site for the G protein may display an atypical behavior in HDX-MS.

**Figure 2.**
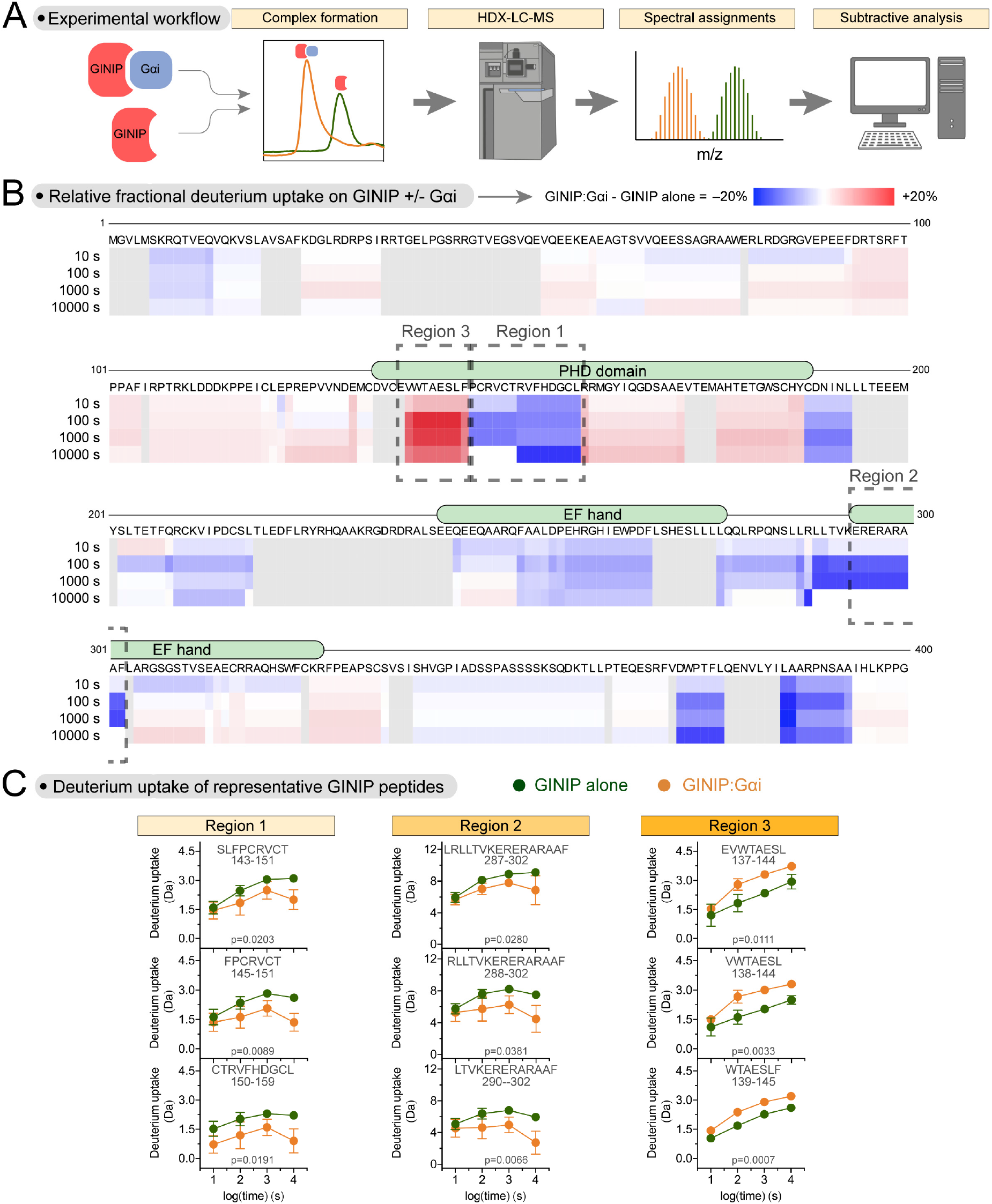
HDX-MS reveals altered protein dynamics in distinct regions of GINIP upon binding Gαi. **(A)** Schematic of HDX-MS workflow. Concentrated protein samples of GINIP alone or in complex with Gαi3 were subjected to hydrogen deuterium exchange (HDX) for different times before in line digestion of peptides that were analyzed by LC-MS to calculate the uptake of deuterium per peptide. **(B)** Gαi3 induces both increases and decreases in deuterium uptake across different regions of GINIP. Stacked heat map of the difference in relative fractional uptake of deuterium by GINIP peptides from the GINIP:Gαi complex relative to GINIP alone at different times (10 s, 100 s, 1000 s, 10000 s), where blue is a decrease in deuterium uptake and red is an increase. White means no difference and grey indicates regions without peptide coverage. Data are the mean differential uptake for an experiment run as a quadruplicate. Boxes indicate the regions with the largest decrease in deuterium uptake (Regions 1 and 2) or the largest increase in deuterium uptake (Region 3). **(C)** Deuterium uptake for GINIP alone or GINIP:Gαi3 at different time points for representative peptides of each one of the three GINIP regions boxed in (B). Deuterium uptake was plotted versus exposure time for three representative peptides of each Region (1-3). Mean ± SD of 4 replicates per time point, except for some of the peptides at 10000s, for which only two to three peptides could be analyzed. p-values reported two-way ANOVA for GINIP alone/GINIP:Gαi x time.

### The first loop of the PHD domain of GINIP is required for binding Gαi3

Prompted by the puzzling results above, we turned our attention to regions of GINIP with *increased* deuterium uptake upon Gαi3 binding. Although atypical, we reasoned that a large increase in local dynamics might indicate a region of interaction if the dynamics for that region of GINIP were somehow constrained in the absence of Gαi3. Only one region in GINIP displayed a marked increase in deuterium uptake (**Fig. 2B, C**), which corresponded to the first loop of the PHD domain. To test if this loop is involved in G protein binding, we generated a chimeric protein (GINIP L1chi) in which this region was replaced by the sequence corresponding to the first loop of another PHD domain (from PHF14, **Fig. 3A**). We found that, in contrast to the wild-type (WT) protein, GINIP L1chi from mammalian cell lysates or purified from bacteria did not bind to purified active Gαi3 (**Fig. 3B, C**). Purified GINIP L1chi had the same thermal stability as GINIP WT (**Fig. S4A**), suggesting that the lack of G protein binding was not a non-specific consequence of overt misfolding of the mutant. Next, we mutated one by one the nine amino acids of this loop, and tested binding of each mutant GINIP expressed in mammalian cells to Gαi3. We found that many of these mutants displayed reduced binding to active Gαi3, including two mutations (V138A and W139A) that completely ablated binding (**Fig. 3D**). We confirmed this lack of binding for GINIP V138A and GINIP W139A with purified proteins (**Fig. 3E**). Purified GINIP V138A and GINIP W139A had the same thermal stability as GINIP WT (**Fig. S4B**), suggesting that these mutations specifically affect G protein binding rather than having a non-specific effect on GINIP’s folding. Finally, we tested the effect of the W139A mutation on the interaction between GINIP and Gαi3 in cells upon GPCR-mediated G protein activation. For this, we generated constructs to detect the interaction between GINIP and Gαi3 using bioluminescence resonance energy transfer (BRET). Two reciprocal donor-acceptor pairs were generated by fusing Gαi3 to nanoluciferase (Nluc) and GINIP to a YFP, or *vice versa* (**Fig. 3F**). When these BRET pairs were co-expressed in HEK293T cells with the GABA_B_ receptor (GABA_B_R), stimulation with GABA led to an increase in BRET that reverted to baseline upon application of a GABA_B_R antagonist, indicating association between GINIP WT and GPCR-activated Gαi3 (**Fig. 3F**). In contrast, introducing the W139A mutation in GINIP nearly abolished the BRET response in this assay (**Fig. 3F**), indicating that the mutation disrupts the GINIP binding to the G protein in cells. Taken together, these results indicate that the first loop of the PHD domain of GINIP contains amino acids that are essential for binding to active Gαi.

**Figure 3.**
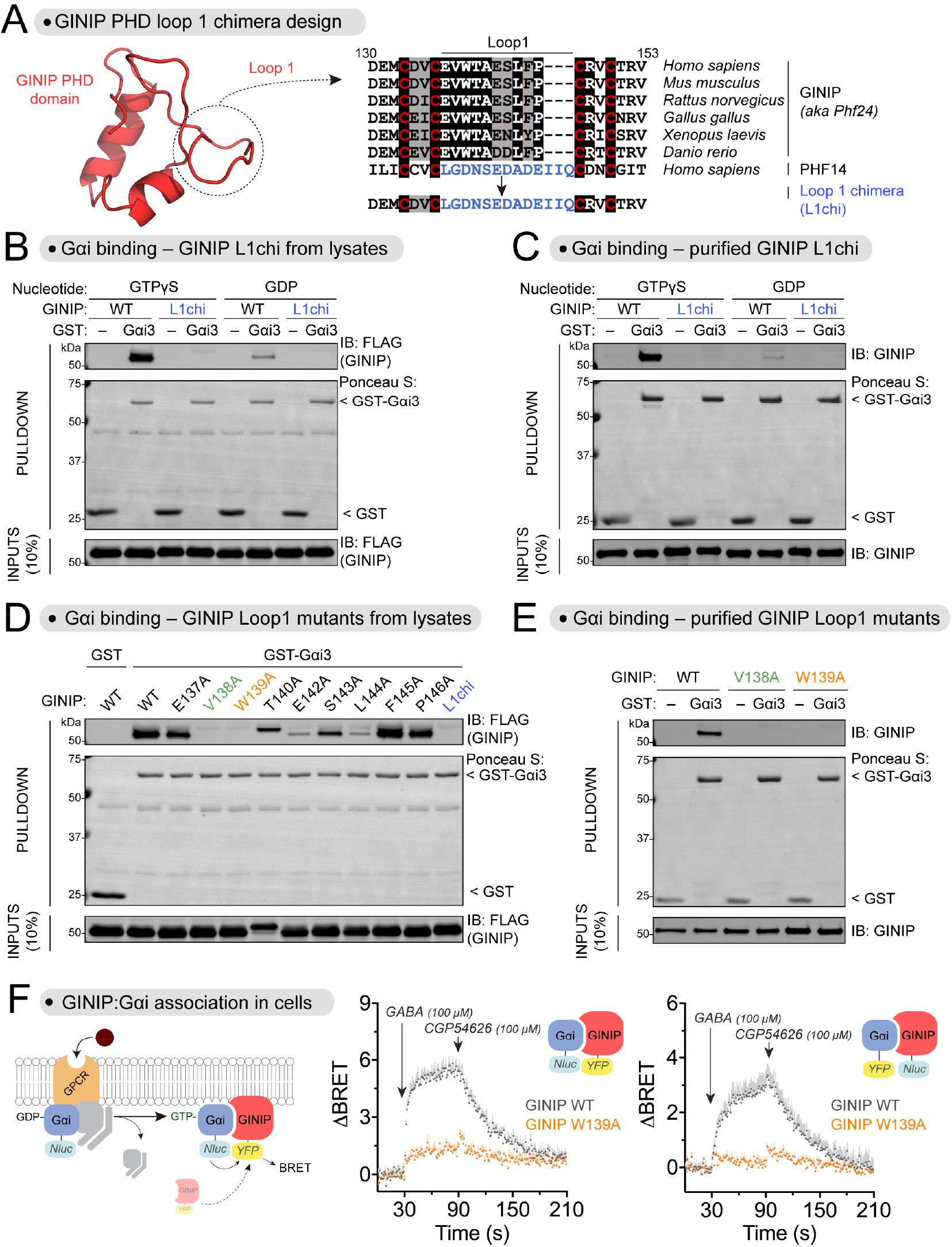
Loop 1 of the PHD domain of GINIP is required for binding Gαi. **(A)** Design of a GINIP chimeric protein construct replacing first loop of the PHD domain of GINIP (Loop 1) with the Loop 1 of the PHD domain of PHF14. *Left,* AlphaFold 2.0 structure of the PHD domain of GINIP (Uniprot #Q9UPV7). *Right,* alignment of sequences corresponding to the Loop 1 of GINIP from different species and the Loop 1 of human PHF14 (in blue). Background shading of the alignment was black or grey if the residue was identical or similar, respectively, in ≥50% of the sequences. Red letters highlight PHD domain conserved cysteine residues at the boundaries of Loop 1. **(B)** GINIP L1chi does not bind to Gαi3-GTPγS. Lysates of HEK293T cells expressing GINIP WT or GINIP L1chi were incubated with GST or GST-Gαi3 immobilized on glutathione-agarose beads in the presence of GDP or GTPγS, as indicated. Bead-bound proteins were detected by Ponceau S staining or by immunoblotting (IB). **(C)** Purified GINIP L1chi does not bind to Gαi3-GTPγS. Purified His-tagged GINIP and GINIP L1chi were incubated with GST or GST-Gαi3 immobilized on glutathione-agarose beads in the presence of GDP or GTPγS as indicated. Bead-bound proteins were detected by Ponceau S staining or by immunoblotting (IB). **(D)** Mutation of V138 or W139 in GINIP disrupts binding to Gαi. Lysates of HEK293T cells expressing the indicated GINIP mutants were incubated with GST or GST-Gαi3 immobilized on glutathione-agarose beads in the presence of GDP or GTPγS, as indicated. Bead-bound proteins were detected by Ponceau S staining or by immunoblotting (IB). **(E)** Purified GINIP V138A or GINIP W139A does not bind to Gαi3-GTPγS. Purified His-tagged GINIP WT, V138A, or W139A were incubated with GST or GST-Gαi3 immobilized on glutathione-agarose beads in the presence of GDP or GTPγS, as indicated. Bead-bound proteins were detected by Ponceau S staining or by immunoblotting (IB). **(F)** GINIP W139A does not associate with Gαi3-GTP upon GPCR stimulation in cells. *Left,* diagram of GPCR-mediated activation of Gαi and BRET-based detection of association between donor/acceptor tagged GINIP/Gαi. *Center,* BRET was measured in HEK293T cells expressing the GABA_B_R, Gαi3-Nluc, and GINIP-YFP WT (grey) or GINIP-YFP W139A (orange), which were treated with GABA and CGP54626 as indicated. Results are expressed as changes in BRET (ΔBRET) relative to the unstimulated baseline. Mean ± S.E.M., n=4. *Right,* same as in *Center*, but with Gαi3 and GINIP constructs in which the BRET donor and acceptor proteins were swapped. Mean ± S.E.M., n=4. All protein electrophoresis results are representative of n ≥ 3 experiments.

### Protein folding models suggest an effector-like mechanism for the engagement of GINIP with active Gαi

While the results above provide compelling evidence for the requirement of GINIP’s PHD loop 1 for binding Gαi, the increased dynamics of this region upon G protein binding observed in HDX-MS experiments remained counterintuitive. To gain further insight into the features of the GINIP-Gαi interface that could help us rationalize these apparently puzzling observations and gain confidence on the proposed G protein binding site, we leveraged computational folding predictions. For this, we generated a structural model of the GINIP-Gαi complex using ColabFold (Mirdita et al., 2022). The model predicted GINIP’s PHD loop 1 as a prominent region of contact with Gαi (**Fig. 4A**), and detailed inspection of this binding interface revealed a striking consistency with our experimental binding results using GINIP mutants in **Fig. 3D**. For example, the two amino acids that resulted in complete ablation of binding when mutated, V138 and W139 (**Fig. 3D**), made extensive contacts with a hydrophobic groove in Gαi according to the structure model (**Fig. 4A**). Similarly, E142, another amino acid that resulted in a marked decrease in binding when mutated (**Fig. 3D**), also appeared to form a salt bridge with K209 of Gαi in the model (**Fig. 4A**). In contrast, other amino acids in this loop like S143, F145 or P146 that did not make direct contacts with Gαi in the model (**Fig. 4A**) also showed modest or not effect on binding when mutated (**Fig. 3D**). On the G protein side of this interface, the model was also very congruent with previous experimental evidence (Park, 2023). First of all, GINIP’s PHD loop 1 docks on a groove formed between the α3 and the switch II of Gαi, a site previously proposed to be essential for GINIP binding based on G protein mutagenesis data (Park, 2023). For example, mutation of Gαi3’s W211, F215, K248, L249, S252, N256 or K257, all of which are predicted by the model to contact GINIP’s PHD loop 1 (**Fig. 4C**), resulted in loss of binding (Park, 2023), whereas mutation of adjacent amino acids like S206 or G42 not predicted by the model to make direct contact (**Fig. 4C**) had no effect on binding (Park, 2023). Thus, the high congruency between extensive protein-protein binding data after site-directed mutagenesis and the structure model prediction lends confidence to the conclusion that GINIP’s PHD loop 1 is a critical structural determinant for the association with G proteins.

**Figure 4.**
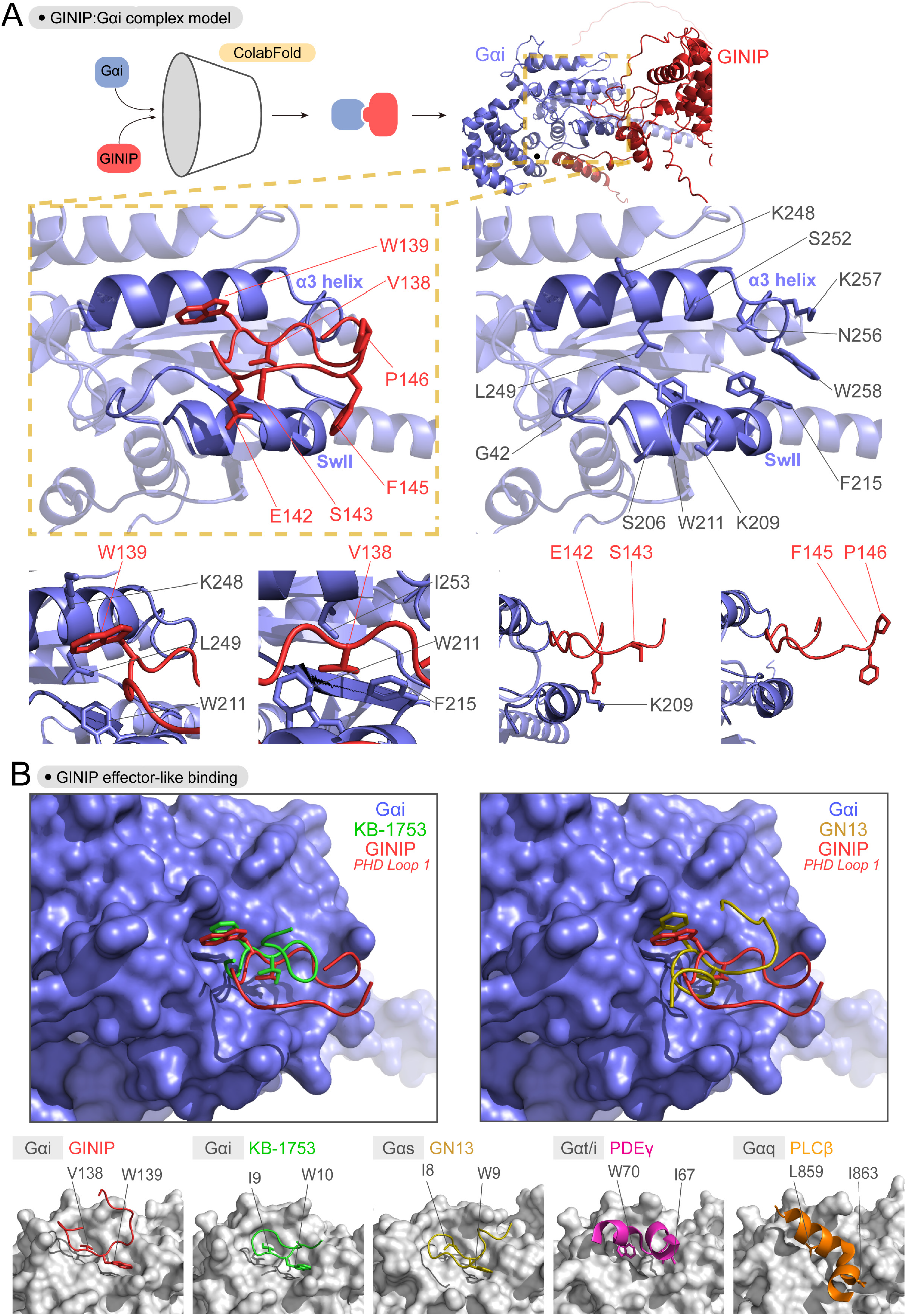
Effector-like binding mode of GINIP Loop 1 on active Gαi. **(A)** Predicted binding pose of Loop 1 of the PHD domain of GINIP onto the α3/SwII groove of Gαi. *Top,* A protein folding model for the complex of GINIP (red) bound to Gαi (blue) was generated using ColabFold. *Middle,* Images depicting the α3/SwII groove of Gαi (blue) and GINIP Loop 1 (red) displaying select side chains. *Bottom,* Close-up views of regions surrounding the indicated GINIP amino acid side chains. **(B)** Comparison of the GINIP:Gαi ColabFold model with structures of Gα subunits in complex with other partners suggests an effector-like binding mode for GINIP. *Left,* Overlay of GINIP’s PHD Loop 1 (red) bound to Gαi (blue) with the Gαi-GTP effector-like peptide KB-1753 (green) . *Right,* Overlay of GINIP’s PHD Loop 1 (red) bound to Gαi (blue) with the Gαs-GTP effector-like peptide GN13 (gold). *Bottom row,* Structural models of GINIP:Gαi1 (red, Colabfold model), KB-1753:Gαi1 (green, PDB ID: 2G83), GN13:Gαs (gold, PDB ID: 7BPH), PDEγ:Gαt/i (pink, PDB ID: 1FQJ), and PLCβ:Gαq (orange, PDB ID: 7SQ2) show conserved positions and orientation of key hydrophobic residues in effector or effector-like partners that mediate G protein binding. Gα subunits are colored grey.

Motivated by the good match between the ColabFold prediction and experimental data, we pursued further evaluation of the features of the GINIP-Gαi interaction according to this model. Binding to the α3/switch II groove Gα is a universal feature of Gα-GTP effectors (Sprang, 1997). While there is no atomic resolution structure of Gαi-GTP bound to an effector, the structure of Gαi1 in complex with the synthetic peptide KB-1753 has been postulated to represent an effector-like binding mode (Johnston et al., 2006). Overlaying this structure with the GINIP-Gαi model revealed not only a similar peptide backbone conformation, but also an almost absolute overlap between the positions of GINIP V138 and W139 with those of KB-1753 I9 and W10, respectively (**Fig. 4B**, which mediate hydrophobic interactions with the α3/switch II groove and are required for KB-1753 binding (Johnston et al., 2006). This high resemblance in binding mode was also shared with another effector-like synthetic peptide for another G protein, Gαs— i.e., the GN13 peptide in complex with Gαs also displays two analogous residues, I8 and W9, in the same positions as V138 and W139 in GINIP or I9 and W10 in KB-1753 (**Fig. 4B**). The same theme of hydrophobic residues docking on the α3/switch II groove has been observed for effectors of other G proteins like Gαt or Gαq (**Fig. 4B**), albeit with some more variation than the examples with peptides above. Overall, these observations support an effector-like binding mode of GINIP on Gαi that is mediated by the first loop of its PHD domain.

### G protein binding induces a long-range conformational change in GINIP

Comparison of the ColabFold model of the GINIP-Gαi complex with the AlphaFold model of GINIP alone provided a potential explanation for the increased dynamics of GINIP’s PHD loop 1 upon Gαi binding seen by HDX-MS. More specifically, we observed that in the model of GINIP alone the first loop of the PHD domain is covered by the N-terminal region of the protein, and that this N-terminal region is displaced upon Gαi binding according to the GINIP-Gαi ColabFold model (**Fig. 5A**). Thus, it is conceivable that the loop 1 of the PHD domain started in a more constrained form in apo-GINIP due to an intramolecular contact, and that once bound to Gαi it displayed increased dynamics. Although appealing, this speculation is based on observations for regions of the models with low confidence prediction scores (**Fig. S5A**), which prompted us to design an experiment to test plausibility. We reasoned that attaching a BRET donor to the N-terminus of GINIP and a BRET acceptor to its C-terminus (Nluc-GINIP-YFP) could report on G protein-induced long range conformational changes. More specifically, we anticipated a decrease in intramolecular BRET due to increased donor to acceptor distance when binding of Gαi displaces the N-terminus of GINIP away from the proximity of the first loop of the PHD domain (**Fig. 5A**). To test this, we carried out a cell-free assay in which purified Gαi3 was added to detergent lysates of cells expressing Nluc-GINIP-YFP (**Fig. 5B**). We found that purified Gαi3-GTPγS caused a dose dependent decrease in BRET with a potency consistent with the affinity of the G protein for GINIP (K_D_∼65 nM) (Park, 2023), whereas GDP-loaded Gαi3, which does not bind to GINIP, had no effect (**Fig. 5B**). We also observed that Gαi3-GTPγS failed to induce a decrease in BRET when the W139A mutation that disrupts G protein binding was introduced in Nluc-GINIP-YFP (**Fig. 5C**), further supporting that the change in GINIP intramolecular BRET was due to Gαi3 binding. To rule out that the decrease in BRET was due to a serendipitous reorientation of the donor and the acceptor unrelated to an increase in distance, we generated three additional constructs in which the position of the donor and the acceptor were swapped and/or in which the position of the N-terminal tag was shifted from the N-terminus to an adjacent region also predicted to move away from the first loop of the PHD domain of GINIP (**Fig. 5A, B**). Albeit with some difference in the amplitude of the BRET changes, all constructs displayed a similar behavior— i.e., a dose dependent decrease in BRET of equivalent potency upon binding of active, GTP-loaded Gαi3 that was not reproduced by inactive, GDP-loaded Gαi3 (**Fig. 5A**) or when the G protein-binding deficient mutant of GINIP W139A was used (**Fig. 5C**). As a complementary approach to test the proposed model, we investigated the effect of truncating the N-terminus of GINIP on Gαi binding. We reasoned that, if GINIP’s N-terminus occludes the first loop of the PHD domain prior to G protein binding, Gαi should bind better in its absence. We found that this is the case because deleting the first 16 or 52 amino acids of GINIP resulted in a modest but reproducible increase in Gαi3 binding (**Fig. S5B**). Taken together with the HDX-MS data, these results support a model in which GINIP undergoes a long range conformational change, likely involving the displacement of a flexible N-terminal region, to accommodate binding of Gαi3 to its PHD domain.

**Figure 5.**
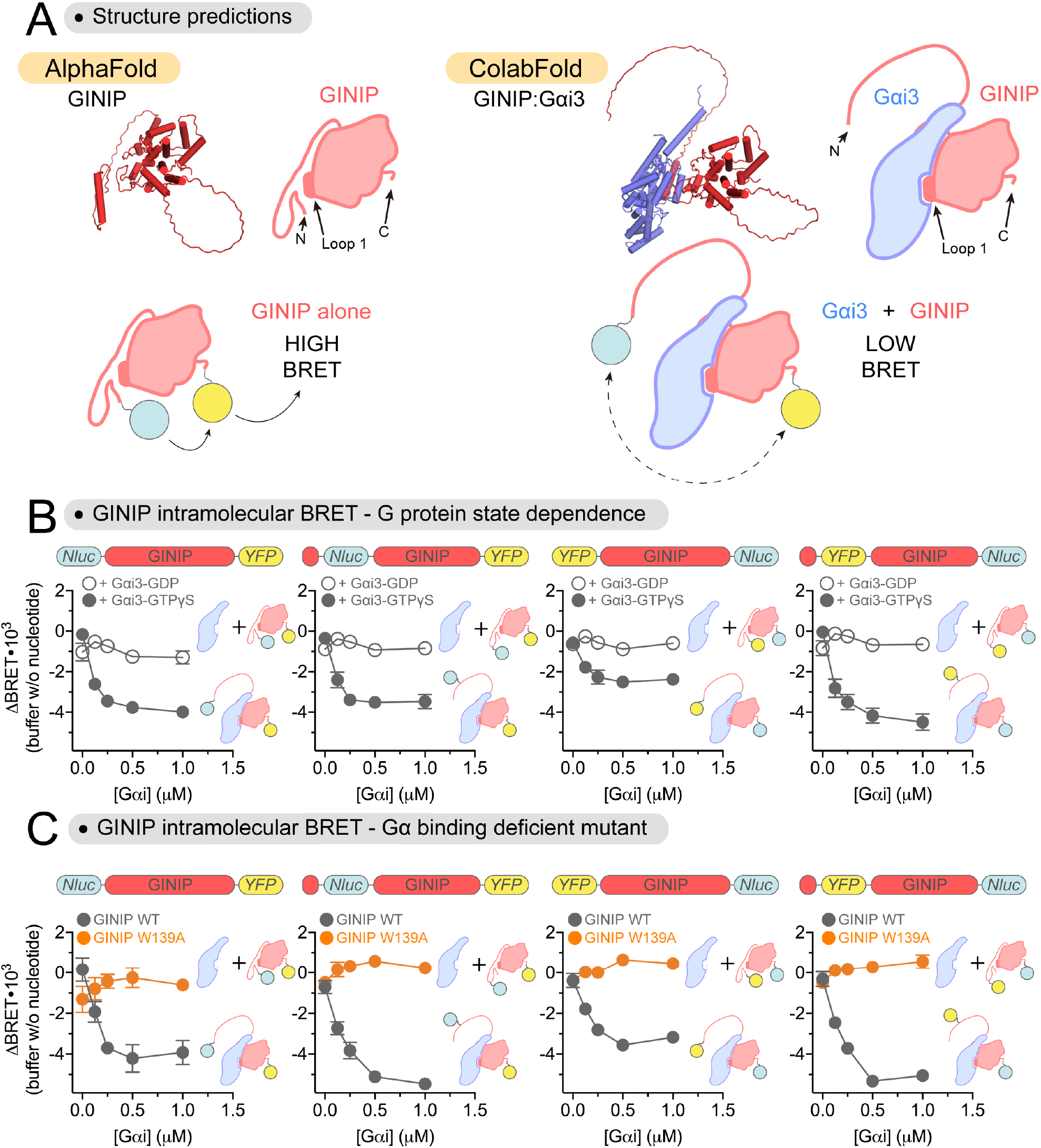
Gαi induces a long-range conformational rearrangement in GINIP. **(A)** *Top*, comparison of structural models of GINIP (red) alone and or in complex with Gαi (blue) suggests displacement of GINIP’s N-termimus from the vicinity of the Loop1 of the PHD domain upon Gαi binding. *Top Bottom,* representative schematic of GINIP intramolecular BRET reporter constructs. Tagging GINIP with both a BRET donor (Nluc, cyan circle) and BRET acceptor (YFP, yellow circle) at the N- and C-terminus could report on G protein-induced long range conformational changes. **(B)** Gαi3-GTPγS induces a dose-dependent decrease in intramolecular BRET for a series of GINIP constructs. Increasing concentrations of purified His-tagged Gαi3 pre-loaded with GDP (open circles) or GTPγS (closed circles) were added to lysates of HEK293T cells expressing the following GINIP intramolecular reporter constructs: Nluc-GINIP-YFP, Nluc(31)-GINIP-YFP, YFP-GINIP-Nluc, and YFP(31)-GINIP-Nluc (represented as bar diagrams above graphs). Mean ± S.E.M., n=3-4. **(C)** Gαi-GTPγS does not induce changes in intramolecular BRET for GINIP constructs bearing the W139A mutation. Increasing concentrations of purified His-tagged Gαi3 pre-loaded with GTPγS were added to lysates of HEK293T cells expressing the same GINIP constructs as in (B), either as WT proteins (gray) or bearing the W139A mutation (orange). Mean ± S.E.M., n=3.

### GINIP’s PHD loop 1 is required to modulate cAMP in cells

Next, we investigated the functional consequences of disrupting the binding of GINIP to G proteins through the PHD loop 1. Previous evidence has established that one consequence of GINIP binding to Gαi-GTP is that it blocks the ability of the G protein to inhibit adenylyl cyclase, thereby dampening the decrease in cellular levels of cAMP observed upon stimulation of GPCRs coupled to Gi (Gaillard et al., 2014; Park, 2023). Consistent with these previous observations, we found that expression of GINIP WT in HEK293T decreased the inhibition of forskolin-induced cAMP upon stimulation of the neurotransmitter receptor GABA_B_R (**Fig. 6A, B**). In contrast, expression of the G protein binding-deficient GINIP mutant V138A or mutant W139A failed to do the same despite expressing at the same levels as GINIP WT (**Fig. 6C**). A similar defect in regulating cAMP responses was observed with expression of the GINIP L1chi construct (**Fig. S6**). in which the G protein binding is disrupted upon replacement of the entire loop 1 of the PHD domain (**Fig. 3A-C**). Meanwhile, another GINIP construct bearing a mutation in an amino acid that is adjacent to V138 and W139 in the PHD loop 1 but that does not affect Gαi binding, F145A (**Fig. 3D**), dampened cAMP inhibition to the same extent as GINIP WT (**Fig. 6B**). These results demonstrate that the first loop of the PHD domain is required for GINIP to modulate cAMP signaling in cells.

**Figure 6.**
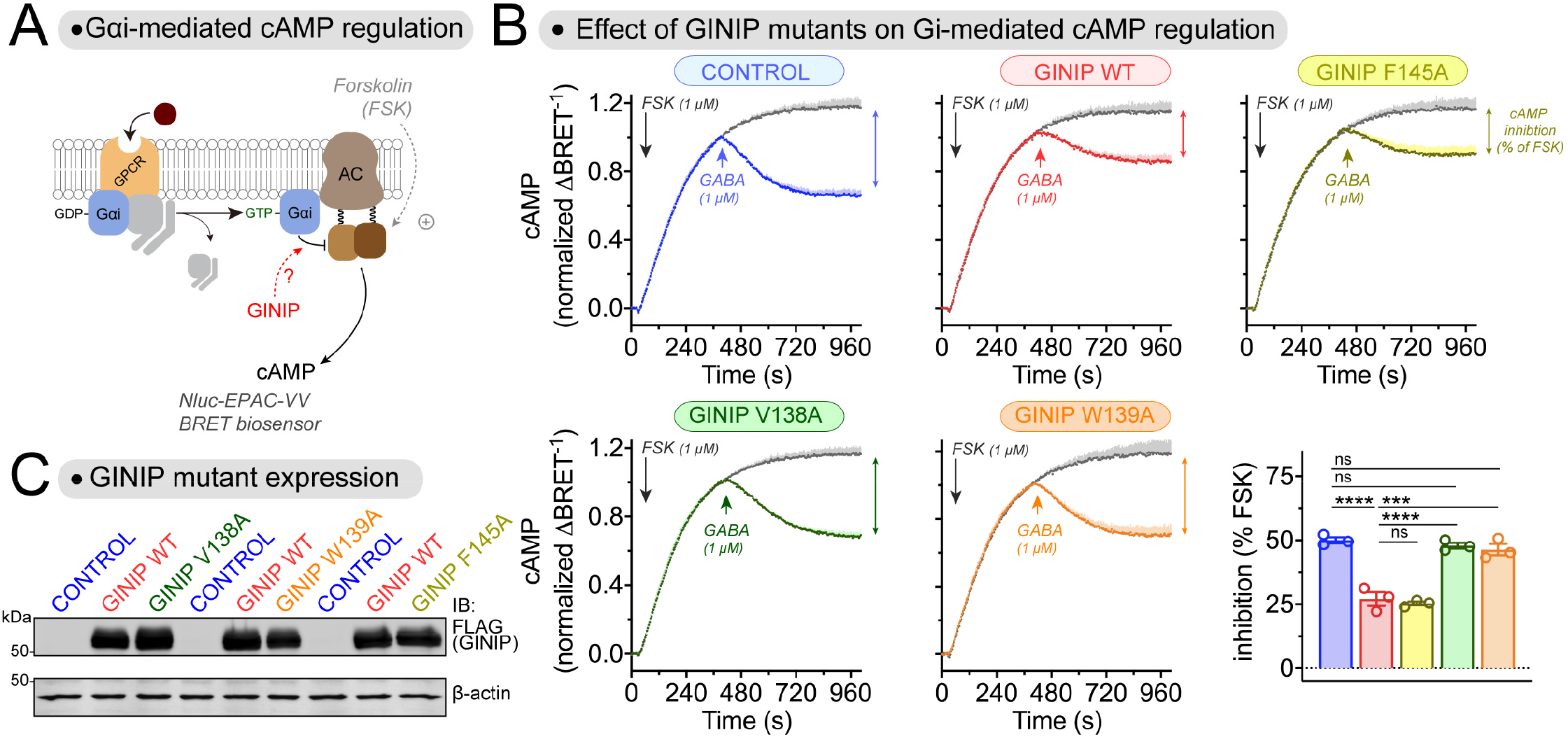
GINIP V138A and GINIP W139A mutants fail to regulate cAMP cellular levels upon GPCR stimulation. **(A)** Diagram of GPCR-mediated activation of Gαi-GTP and subsequent regulation of cAMP levels in cells monitored by BRET. **(B)** Mutation of GINIP V138 or W139, but not F145, prevents the blockade of cAMP inhibition upon stimulation of GABA_B_R observed with GINIP WT. Kinetic BRET measurements of cAMP levels were carried out in HEK293T cells expressing the GABA_B_R without GINIP (blue) or expressing GINIP WT (red), GINIP F145A (yellow), GINIP V138A (green), or GINIP W139A (orange). Cells were treated with forskolin (FSK) and GABA as indicated. Quantification of the inhibition of FSK-stimulated cAMP upon stimulation of GABA_B_R with GABA is shown in the bar graph on the bottom left corner. Mean ± S.E.M., n=3. ns = not significant, ***p<0.001, ****p<0.0001, one-way ANOVA corrected for multiple comparisons (Tukey). **(C)** Representative immunoblotting (IB) result confirming equal expression of GINIP WT, GINIP V138A, GINIP W139A, and GINIP F145A in the cells used for the experiments shown in (B).

### GINIP’s PHD loop 1 is required to regulate free Gβγ levels in cells

Another previously reported consequence of GINIP on G protein signaling is that it favors free Gβγ signaling by preventing the action of RGS GAPs on Gαi-GTP (Park, 2023). To assess this function, we used a BRET assay that monitors the levels of free Gβγ in HEK293T cells (**Fig. 7A**). One effect of GINIP in this system is that it slows deactivation rates when GPCR signaling is shut off with an antagonist after agonist stimulation, reflecting a longer lifetime of Gβγ in a free state before reassociation with Gα (Park, 2023). While GINIP WT slowed deactivation of free Gβγ upon GABA_B_R modulation, GINIP V139A or GINIP W139A failed to do so (**Fig. 7A**). GINIP WT also reverts the effects of RGS protein overexpression in this system (Park, 2023), which are a reduction of Gβγ response amplitudes upon GPCR agonist stimulation and an acceleration of Gβγ deactivation rates (Park, 2023). For example, expression of RGS8 decreases Gβγ responses to GABA_B_R stimulation and accelerates deactivation (**Fig. 7B**). While co-expression of GINIP WT restores response amplitudes and deactivation rates to levels equivalent to those observed in controls, co-expression of GINIP V138A or GINIP W139A did not (**Fig. 7B**). Similar observations were made with RGS12 (**Fig. 7C**), which belongs to an RGS family different from that of RGS8. Also, like with GINIP V138A and GINIP W139A, expression of GINIP L1chi failed to recapitulate the effects of GINIP WT in any of the aspects of free Gβγ signaling regulation investigated (**Fig. S7**). These results indicate that the first loop of the PHD domain is required not only for GINIP to modulate cAMP responses, but also Gβγ signaling in response to GPCR stimulation.

**Figure 7.**
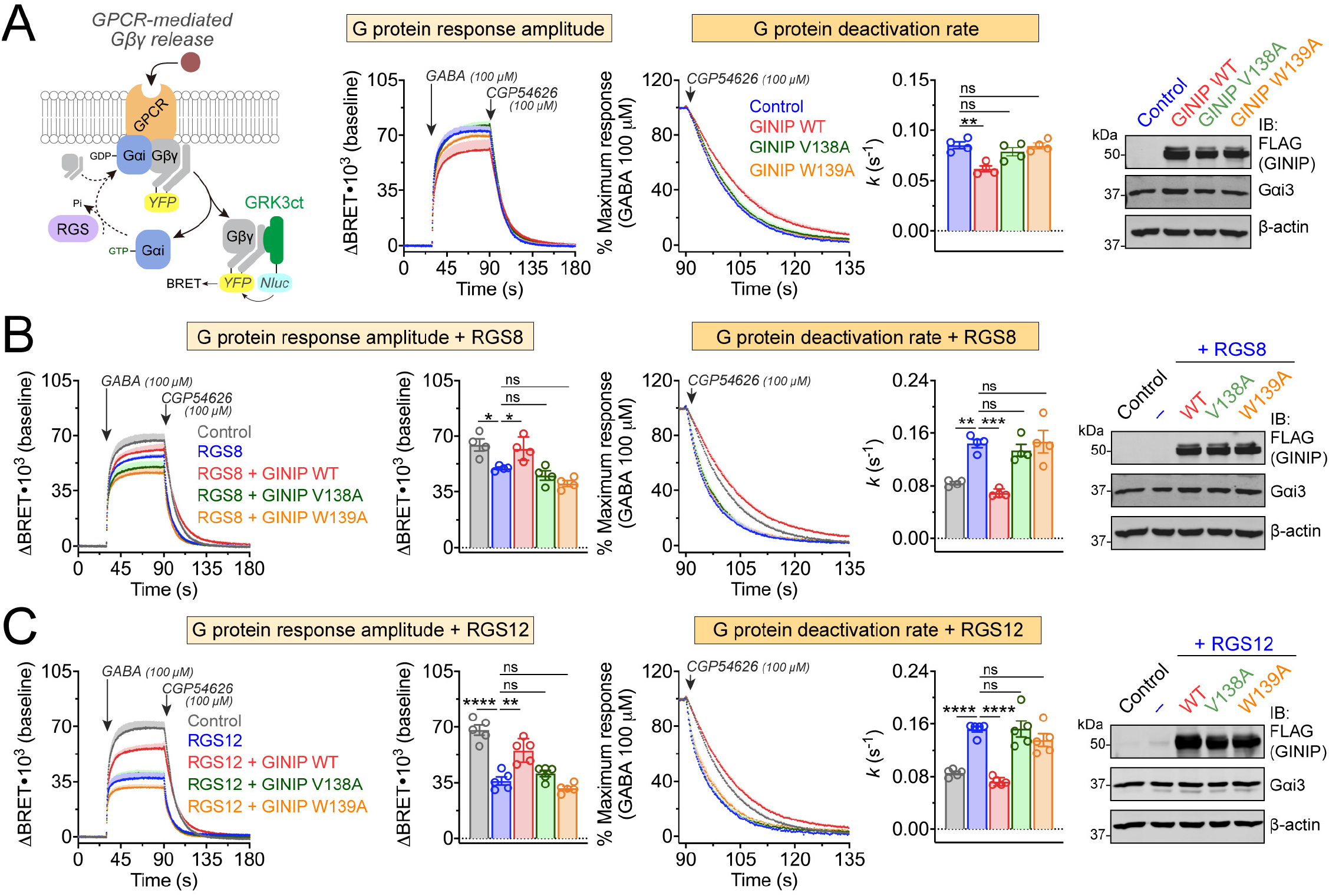
GINIP V138A and GINIP W139A mutants fail to regulate Gβγ responses in cells upon GPCR stimulation. **(A)** Mutation of GINIP V138 or W139 prevents the enhancement of Gβγ signaling upon stimulation of GABA_B_R observed with GINIP WT. *Left,* diagram of G protein activation/deactivation cycle and BRET-based detection of free Gβγ. *Center,* kinetic BRET measurements were carried out in HEK293T cells expressing the GABA_B_R without GINIP (blue), or expressing GINIP WT (red), GINIP V138A (green), or GINIP W139A (orange). Cells were treated with GABA and CGP54626 as indicated. *Right,* G protein deactivation rates were determined by normalizing the BRET data to maximum response and fitting the post-antagonist data to an exponential decay curve to extract rate constant values (*k*). Mean ± S.E.M., n=4. ns = not significant, **p<0.01, one-way ANOVA corrected for multiple comparisons (Tukey). A representative immunoblotting (IB) result confirming equal expression of GINIP WT or mutants, and Gαi3 in these experiments is shown on the right. **(B)** Mutation of GINIP V138 or W139 prevents the RGS8-mediated regulation of Gβγ signaling upon stimulation of GABA_B_R observed with GINIP WT. BRET experiments were carried out and analyzed as in (A) with cells expressing RGS8 alone (blue), or RGS plus GINIP WT (red), GINIP V138A (green), or GINIP W139A (orange), or neither RGS8 nor GINIP (grey). Quantification of G protein response amplitude was determined 1 minute after agonist stimulation. Mean ± S.E.M., n=4. ns = not significant, *p<0.05, **p<0.01, ***p<0.001, one-way ANOVA corrected for multiple comparisons (Tukey). A representative immunoblotting (IB) result confirming equal expression of GINIP WT or mutants, and Gαi3 in these experiments is shown on the right. **(C)** Mutation of GINIP V138 or W139 prevents the RGS12-mediated regulation of Gβγ signaling upon stimulation of GABA_B_R observed with GINIP WT. BRET experiments were carried out and analyzed as in (A) with cells expressing RGS12 alone (blue), or RGS plus GINIP WT (red), GINIP V138A (green), or GINIP W139A (orange), or neither RGS12 nor GINIP (grey). Quantification of G protein response amplitude was determined 1 minute after agonist stimulation. Mean ± S.E.M., n=5. ns = not significant, **p<0.01, ****p<0.0001, one-way ANOVA corrected for multiple comparisons (Tukey). A representative immunoblotting (IB) result confirming equal expression of GINIP WT or mutants, and Gαi3 in these experiments is shown on the right.

## DISCUSSION

The main advance provided by this work is the identification of a structural element of GINIP required for its interaction with Gαi and that the binding mediated by this element determines the modulation of neurotransmitter GPCR signaling by GINIP. More specifically, our results show that GPCR signaling modulation by GINIP requires the engagement between the loop 1 of its PHD domain and the α3/SwII groove of Gαi, a classical effector-binding region for Gα proteins. The functional consequence of this interaction on GPCR signaling is a simultaneous dampening of Gαi-mediated signaling and enhancement of Gβγ-mediated signaling (Park, 2023). By strongly supporting an effector-like engagement of the PHD loop 1 with Gαi, our results sheds further light onto the mechanism by which GINIP regulates GPCR signaling. On one hand, this effector-like binding mode is compatible with competition between GINIP and its natural effector adenylyl cyclase for binding to the α3/SwII groove on Gαi, which explains the dampening cAMP modulation exerted by GINIP. On the other hand, this binding mode would also be expected to prevent RGS protein binding by steric hindrance with the bulk of the PHD domain, which explains the enhancement of free Gβγ signals. Given that the effects of GINIP on G protein signaling have been validated with different downstream readouts, like cAMP for Gαi or GIRK channel activity for free Gβγ, in physiologically-relevant systems, like neurons (Park, 2023), the mechanistic insights gained here are broadly applicable to GINIP’s roles in controlling neurological processes (Gaillard et al., 2014; Kuramoto et al., 2017; Park, 2023).

Our identification of the Gαi binding site on GINIP as a region of increased deuterium exchange was initially surprising because ligand binding often results in decreases in deuterium uptake. Although we can only speculate about the structural basis that would explain this, the evidence to support the requirement of the loop 1 of the PHD domain for G protein binding is very strong. As for what could account the puzzling behavior in HDX-MS, we provide a plausible explanation by combining several independent lines of evidence. First, we gained confidence on the protein complex prediction by cross validation with experimental protein-protein binding data (**Fig. 3D**). Then, the suggestion from folding predictions that the N-terminus might initially constrain the loop 1 of PHD domain of GINIP (**Fig. S5A**), thereby providing a potential explanation for its increased dynamics upon G protein binding, was corroborated by BRET-based (**Fig. 5**) and protein binding experiments (**Fig. S5B**). Although it is tempting to believe our model that GINIP undergoes a long range conformational change to accommodate the engagement of Gαi to the PHD domain, atomic resolution structural information will be required in the future to provide definitive information.

To our knowledge, the results presented here are the first example of an annotated PHD domain with a function different from epigenetic or transcriptional regulation (Li et al., 2006; Peña et al., 2006). PHD domains are frequently found in proteins that recognize modified and unmodified histone tails to participate in the regulation of chromatin condensation or, less frequently, in proteins involved in transcriptional regulation (Fiedler et al., 2008; Miller et al., 2010) that bind to histone modifications and DNA. In contrast, our results indicate that the PHD domain of GINIP has a non-nuclear, signaling-related function that directly controls the processing of receptor initiated responses (Musselman and Kutateladze, 2011; Sanchez and Zhou, 2011). Since PHD domains in the proteome have not been thoroughly characterized, it is possible that other examples of atypical functions, including, but not limited to, G protein regulation, could emerge in the future. Since PHD domains have been proposed to be druggable targets (Amato et al., 2018), our results presented here raise the possibility of GINIP as a new actionable point within GPCR signaling.

The mechanism by which GINIP modulates GPCR signaling, i.e., by enhancing the amplitude and duration of Gβγ-mediated signaling to the detriment of Gαi-mediated signaling, is still not known to be shared with any other protein. However, the identification of a structural element essential for GINIP binding to G proteins sets the basis to identify other regulators of the same class by sequence and structural similarity. Although we have not carried out a systematic search for potential GINIP-like G protein regulators, we were intrigued by the small family of proteins named G protein-regulated inducer of neurite outgrowth 1, 2 and 3 (GRIN1, GRIN2 and GRIN3). Much like GINIP, GRIN proteins bind to GTP-bound Gα subunits of the G_i/o_ family (Iida and Kozasa, 2004; Mototani et al., 2018; Nakata and Kozasa, 2005); GRIN1 is reported to have no effect on the ability of Gα subunits to bind or hydrolyze nucleotides (Nakata and Kozasa, 2005) and their functional interplay with G proteins remains poorly understood. Using a folding model of GRIN3 with Gαi1 (**Fig. S8**), we noticed a striking similarity between the predicted binding of GRIN3 and that observed for GINIP or the effector-like peptide KB-1753. More specifically, the two hydrophobic residues V695 and Y696 of GRIN3 dock on the α3/swII groove in the same fashion as GINIP’s V138 and W139 (**Fig. S8**). These amino acids are in a previously identified G protein binding region in GRIN proteins (Nakata and Kozasa, 2005). Moreover, the amino acid region presumably used by GRIN3 to engage Gα is not only conserved across all GRIN proteins, but also shares similarity with GINIP’s PHD loop 1. Sequence alignment suggests a potential short consensus motif [–]-Ψ-W-x-π, where [–] is a negatively charged residue, Ψ is a residue with an aliphatic chain, W is an aromatic residue, x is any residue, and π is a residue with a small side-chain. Intriguingly, Nakata and Kozasa reported, as data not shown, that “*The inhibitory effect of GRIN1 on the GAP activity of RGS4 for Gαo indicates the competition of the interaction of RGS4 and GRIN1 on Gαo*” (Nakata and Kozasa, 2005), suggesting further functional similarities between GINIP and GRIN proteins. While we describe the similarities between GINIP and GRIN proteins as speculation, we believe that the comparison exemplifies how the information gained here about the molecular basis for GINIP-mediated G protein regulation could be leveraged to generate hypotheses about putative G protein regulators with GINIP-like properties.

In summary, this work sheds further light onto the molecular basis for mechanisms that regulate neuromodulatory GPCR responses by acting at the immediate post-receptor level. These post-receptor events are still poorly defined, and, while the functions of GINIP itself have important physiological implications in the nervous systems (Gaillard et al., 2014; Kuramoto et al., 2017; Serikawa et al., 2019), the findings reported here could be leveraged to gain deeper understanding of GPCR and G proteins signaling regulation in other contexts.

## EXPERIMENTAL PROCEDURES

### Plasmids

The plasmid for the bacterial expression of C-terminally His-tagged human GINIP (pET21a(+)-GINIP-His) was generated by removing the ligation independent cloning (LIC) cassette from a previously described vector, pLIC-His (Stols et al., 2002), using NdeI and HindIII and inserting by Gibson assembly the sequence of human GINIP (NCBI Gene ID: 23349) followed by a TEV cleavage site upstream of the C-terminal His-tag that remained in the pLIC-His plasmid after digestion. Removal of the LIC cassette reverted the plasmid back to the parental vector, pET21a(+), and was renamed as such. The same approach was followed to generate GINIP truncations, ΔN16 and ΔN53, as well as the GINIP L1chi constructs in which Loop 1 of the PHD domain of GINIP (aa 137-146) was replaced by the sequence of the Loop 1 of the PHD domain of human PHF14 (aa 326-338; NCBI Gene ID: 9678). To generate GINIP protein used in HDX-MS experiments, a second bacterial expression plasmid was generated using a human GINIP sequence codon optimized for expression in *E. coli*, which was inserted into pET21a(+) without a TEV cleavage site preceding the His-tag (pET21a(+)-coGINIP-His). Plasmids for the bacterial expression of rat His-Gαi3 (pET28b-Gαi3) and GST-Gαi3 (pGEX-4T-1-Gαi3) have been described previously (Garcia-Marcos et al., 2009; Garcia-Marcos et al., 2020; Marivin et al., 2020). The pbb131 plasmid encoding yeast *N*-myristoyltransferase (NMT) (Mumby and Linder, 1994) was a gift from Maurine Linder (Cornell University).

The plasmid for mammalian expression of C-terminally 3xFLAG-tagged GINIP (p3xFLAG-CMV-14-GINIP) was made by digesting p3xFLAG-CMV-14 with EcoRI and BamHI and inserting the human GINIP sequence upstream of the FLAG-tag (p3xFLAG-CMV-14-GINIP). The plasmid encoding the chimeric GINIP L1chi construct (p3xFLAG-CMV-14-GINIPL1chi) was made similarly, except that a multi-fragment Gibson assembly strategy was implemented to insert the sequence of PHF14 described above. p3xFLAG-CMV-14-GINIP was also used as the starting point to generate the intramolecular BRET constructs fused simultaneously to Nluc and YFP described in **Fig. 5**. Nluc or YFP were inserted by Gibson assembly into the BamHI site of p3xFLAG-CMV-14-GINIP to create C-terminal fusions to GINIP. Then, Nluc or YFP were fused to the N-terminus of GINIP by insertion into the EcoRI site of the same constructs, or to an internal position between amino acids 31 and 32 of GINIP by insertion into an StuI site. In all cases, GGGS linker sequence was introduced between GINIP and Nluc or YFP. The four resulting plasmids were p3xFLAG-CMV-14-Nluc-GINIP-YFP, p3xFLAG-CMV-14-Nluc(31)-GINIP-YFP, p3xFLAG-CMV-14-YFP-GINIP-Nluc, and p3xFLAG-CMV-14-YFP(31)-GINIP-Nluc. The plasmid encoding untagged Gαi3 (pcDNA3.1(+)-Gαi3 WT) was described previously (Garcia-Marcos et al., 2010). pcDNA3.1(+)-GABA_B_R1a and pcDNA3.1(+)-GABA_B_R2 were a gift from Paul Slessinger (Icahn School of Medicine at Mount Sinai, NY). Plasmids encoding mas-GRK3ct (pcDNA3.1-masGRK3ct-Nluc) and the cAMP sensor Nluc-EPAV-VV (pcDNA3.1-Nluc-EPAC-VV) were provided by K. Martemyanov (Scripps Research Institute, FL) (Masuho et al., 2015). pcDNA3.1-Venus(1–155)-Gγ_2_ (VN-Gγ2), and pcDNA3.1-Venus(155-239)-Gβ1 (VC-Gβ1) were a gift from N. Lambert (Augusta University, GA) (Hollins et al., 2009). pcDNA3.1(-)-3xHA-RGS8 was acquired from the cDNA Resource Center (RGS080TN00) (Bloomsberg University, PA). The plasmids encoding rat Gαi3 tagged with Nluc or YFP in the a/b loop, pcDNA3.1(-)-Gαi3-Nluc(a/b) or pcDNA3.1(-)-Gαi3-YFP(a/b), respectively, were generated by inserting EcoRI and XhoI restriction sites between residues 91 and 92 of Gαi3, and then inserting Nluc or YFP by Gibson assembly at those sites, which were maintained in the final construct. The resulting construct contains a 5’ linker encoding the amino acids Glu-Phe and 3’ linker encoding the amino acids Ser-Ser flanking the Nluc or YFP insert sequence. The plasmid encoding for mouse RGS12 (pCMV-Sport6-RGS12) was obtained from the DNA Resource Core PlasmID Repository (Harvard Medical School, Plasmid ID MmCD00316157). All point mutations were generated using QuikChange II following the manufacturer’s instructions (Agilent, 200523).

### Protein expression and purification

His-tagged and GST-tagged Gαi3 were expressed in BL21(DE3) *E. coli* transformed with the corresponding plasmids by overnight induction at 23 °C with 1 mM isopropyl β-D-1-thio-galactopyranoside (IPTG). His-tagged GINIP was induced for 5 hours at 23 °C with 1 mM IPTG. In all cases, IPTG was added when the OD_600_ reached ∼0.8. Unless otherwise indicated, protein purification was carried out following previously described protocols (Garcia-Marcos et al., 2009; Marivin et al., 2020). Briefly, bacteria pelleted from 1 liter of culture were resuspended at 4 °C in 25 ml of lysis buffer (50 mM NaH_2_PO_4_, pH 7.4, 300 mM NaCl, 10 mM imidazole, 1% (v/v) Triton X-100, supplemented with a protease inhibitor mixture of 1 μM leupeptin, 2.5 μM pepstatin, 0.2 μM aprotinin, and 1 mM phenylmethylsulfonyl fluoride). When purifying Gαi3, this buffer was supplemented with 25 μM GDP and 5 mM MgCl_2_. After sonication (4 pulses of 30 s separated by 30 s intervals for cooling), the lysate was cleared by centrifugation at 12,000 × *g* for 30 min at 4 °C. The soluble fraction was used for affinity purification in batch on HisPur Cobalt (Thermo, 89964) or GSH-agarose resins (Thermo, 16100) by incubating lysate and beads with rotation for 2 hours at 4 °C. Resin was washed 3 times with lysis buffer and then eluted with lysis buffer supplemented with 250 mM imidazole or with 50 mM Tris-HCl, pH 8, 100 mM NaCl, 30 mM reduced GSH, respectively. His-tagged GINIP proteins were buffer exchanged to PBS (137 mM NaCl, 2.7 mM KCl, 8 mM Na_2_HPO_4_, and 2 mM KH_2_PO_4_) by overnight dialysis (10,000 Da cut-off) at 4 °C with the exception of protein used for HDX-MS (described in detail below). His-Gαi3 and GST-Gαi3 were buffer exchanged to 20 mM Tris-HCl, pH 7.4, 20 mM NaCl, 1 mM MgCl_2_, 1 mM DTT, 10 μM GDP, 5% (v/v) glycerol using a HiTrap desalting column (Cytiva, 29048684) connected to an AKTA FPLC.

For HDX-MS experiments, purified His-Gαi3 protein was concentrated by centrifugation at 4,300 x g with Amicon Ultra 4 10,000 Da NMWL centrifugal filter (MilliporeSigma, UFC801024) to 315 μM after purification. The concentrated Gαi3 stock was then supplemented with 3 mM GTPγS and incubated at 30 °C for 3 hours to allow for nucleotide loading before aliquoting and storing at −80 °C. For the GINIP protein produced for HDX-MS experiments, the eluate from the HisPur Cobalt resin was subjected to ion-exchange chromatography in a HiTrapQ HP column (Cytiva, 17115401). GINIP-containing fractions were buffer exchange to HDX-MS buffer (20 mM HEPES pH 7.4, 150 mM NaCl, 10 mM MgCl_2_, 1 mM DTT) using a HiTrap desalting column and then concentrated to 260 μM using Amicon Ultra 4 10,000 Da NMWL centrifugal filter before aliquoting and storing at −80 °C.

Myristoylated GINIP (myr-GINIP) was purified from BL21(DE3) *E. coli* bacteria co-expressing the plasmid encoding pET21a(+)-GINIP-His with a plasmid encoding N-myristoyl transferase (NMT) as described above for non-myrisroylated GINIP, except that the eluate from the HisPur Cobalt resin was subjected to ion-exchange chromatography in a HiTrapQ HP column (Cytiva, 17115401).

### Gel filtration chromatography

For experiments shown if **Fig. 1A**, GINIP and Gαi3-GTPγS concentrated protein stocks were diluted separately or together in HDX-MS buffer at a concentration of 5 μM and 5.6 μM, respectively. After 15 minutes of equilibration in ice, 235 μl of the samples were injected into a Superdex 200 Increase 30/100 GL column connected to an AKTA FPLC kept at 4 °C and pre-equilibrated with HDX-MS buffer. For experiments shown if **Fig. 1B**, GINIP and Gαi3-GTPγS concentrated protein stocks were diluted separately or together in HDX-MS buffer at a concentration of 58.9 μM and 118 μM, respectively, and incubated for 15 minutes (t=0) or 24 h (t=24h). Twenty μl of each sample were diluted 14-fold in HDX-MS buffer made with 99.9% D_2_O (151882-100G, Sigma-Aldrich) to mimic the exposure conditions for HDX-MS experiments, and the entire volume was injected into a Superdex 200 Increase 30/100 GL column connected to an AKTA FPLC kept at 4 °C. Column was run with a 1.5 column volumes of isocratic elution with HDX-MS buffer and 0.5 ml fractions were collected. Thirty-two μl of each fraction were mixed with Laemmli sample buffer and incubated at 65 °C for 10 min before separation of proteins by SDS-PAGE and staining with Coomassie blue. Chromatograms were normalized to the maximum intensity of the signal (mAU) detected for each series of samples.

### Pulldown assays

For experiments using purified proteins as source of soluble binding ligands, GST or GST-Gαi3 were supplemented with 150 μM GDP or GTPγS as indicated in figures and incubated at 30 °C for 3 hours for nucleotide loading. GST-fused proteins were immobilized on GSH-agarose beads (Thermo, 16100) for 90 min at room temperature in 50 mM Tris-HCl, pH 7.4, 100 mM NaCl, 0.4% (v/v) Nonidet P-40, 5 mM EDTA, 2 mM DTT supplemented with 30 μM GDP or 30 μM GTPγS as indicated in the figures. Beads were washed twice with binding buffer and resuspended in 400 μl of binding buffer containing the appropriate nucleotides. 1-2 μg of the following C-terminally His-tagged GINIP proteins were used: GINIP WT, myr-GINIP, GINIP L1chi, GINIP V138A, or GINIP W139A. Aliquots of protein stored at -80 °C were quickly thawed and cleared by centrifugation at 14,000 × *g* for 2 minutes before addition to tubes containing the GST-fused proteins immobilized on GSH-agarose beads in a final volume of 400 μl. Tubes were incubated for 4 h at 4 °C with constant rotation. Beads were washed three times with 1 ml of wash buffer (4.3 mM Na_2_HPO_4_, 1.4 mM KH_2_PO_4_, pH 7.4, 137 mM NaCl, 2.7 mM KCl, 0.1% (v/v) Tween 20, 10 mM MgCl_2_, 5 mM EDTA, 1 mM DTT) supplemented with 30 μM GDP or 30 μM GTPγS, and resin-bound proteins were eluted with Laemmli sample buffer by incubation at 65 °C for 10 min. Proteins were separated by SDS-PAGE and immunoblotted with antibodies as indicated under “*Protein Electrophoresis and Immunoblotting*.”

For experiments using lysates of cultured cells as a source of soluble binding ligands, HEK293T cells (ATCC CRL-3216) were grown at 37 °C, 5% CO_2_ in DMEM (Gibco, 11965-092) supplemented with 10% fetal bovine serum (Hyclone, SH30072.03), 100 units/ml penicillin, 100 μg/ml streptomycin, and 2 mM L-glutamine (Corning, 30-009-CI). Approximately 400,000 HEK293T cells were seeded in 6 well plates and transfected the day after using the calcium phosphate method with plasmids encoding the GINIP constructs indicated in the figures. Cell medium was changed 6 h after transfection, and approximately 24 h later, cells were lysed at 4 °C with 120 μl of lysis buffer (20 mM HEPES, pH 7.2, 125 mM K(CH_3_COO), 0.4% (v/v) Triton X-100, 1 mM DTT, 10 mM β-glycerophosphate, and 0.5 mM Na_3_VO_4_ supplemented with a SigmaFAST protease inhibitor mixture (Sigma, S8830)). Cell lysates were cleared by centrifugation at 14,000 × *g* for 10 minutes, and supplemented with 30 μM GDP (GDP condition), or 30 μM GDP, 30 μM AlCl_3_, and 10 mM NaF (GDPꞏAlF ^−^ condition), or 30 μM GTPγS (GTPγS condition) as indicated in the figures. For experiments with GDP and GTPγS conditions, GST or GST-Gαi3 were first supplemented with 150 μM GDP or GTPγS and incubated at 30 °C for 3 hours for nucleotide loading. For experiments with GDPꞏAlF ^−^ conditions, GST or GST-Gαi3 were diluted in binding buffer supplemented with GDPꞏAlF ^−^ after thawing. GST-fused proteins were immobilized on GSH-agarose beads as described above and 100 μl of cell lysates were added to the immobilized GST proteins in a final volume of 400 μl. Tubes were incubated for 4 h at 4 °C with constant rotation, then washed and eluted as described above for purified soluble ligands.

For experiments shown in **Fig. S1C**, GST or GST-Gαi3 were nucleotide loaded with GDP or GTPγS as described above before immobilizing on GSH-agarose beads. His-tagged GINIP proteins were diluted in a modified binding buffer (50 mM Tris-HCl, pH 7.4, 100 mM NaCl, 0.4% (v/v) Nonidet P-40, 100 μM EDTA, 2 mM DTT) for stripping of divalent cations by EDTA, or in the same buffer without EDTA for controls. 15 μl of diluted GINIP proteins were then added to GSH-agarose bound GST proteins resuspended in 285 μl of binding buffer supplemented or not with 200 μM CaCl_2_ or 200 μM ZnSO_4_, as indicated in the figure, as well as 30 μM GDP or 30 μM GTPγS. Tubes were incubated for 4 h at 4 °C with constant rotation, then washed three times with 1 ml of a modified wash buffer (4.3 mM Na_2_HPO_4_, 1.4 mM KH_2_PO_4_, pH 7.4, 137 mM NaCl, 2.7 mM KCl, 0.1% (v/v) Tween 20, 10 mM MgCl_2_, 100 μM EDTA, 1 mM DTT) supplemented with 30 μM GTPγS and 200 μM CaCl_2_ or 200 μM ZnSO_4_ as indicated in the figure. Samples were eluted as described above for purified soluble ligands.

### Protein electrophoresis and immunoblotting

Protein samples were prepared in Laemmli sample buffer as described in other sections. Proteins were separated by SDS-PAGE and transferred to PVDF membranes, which were blocked with 5% (w/v) nonfat dry milk and sequentially incubated with primary and secondary antibodies diluted in 2.5% (w/v) nonfat dry milk with 0.05% (w/v) sodium azide. For protein-protein binding experiments with GST-fused proteins, PVDF membranes were stained with Ponceau S and scanned before blocking. The primary antibodies used were the following (dilution factor in parenthesis): rabbit GINIP, Aviva ARP70657_P050 (1:2000); rabbit Gαi3, SCBT #sc-262 (1:1000); mouse FLAG, Sigma #F1804 (1:1000); rabbit β-actin, LI-COR #926-42212 (1:1000). The secondary antibodies were (dilution factor in parenthesis): goat anti-rabbit Alexa Fluor 680, Invitrogen #A21077 (1:10,000); goat anti-mouse Alexa Fluor 680, Invitrogen #A21058 (1:10,000); goat anti-mouse IRDye 800, LI-COR #926-32210 (1:10,000); goat anti-rabbit DyLight 800, Thermo #35571 (1:10,000). Infrared imaging of immunoblots was performed using an Odyssey CLx Infrared Imaging System (LI-COR). Images were processed using ImageJ software (National Institutes of Health) or Image Studio software (LI-COR), and assembled for presentation using Photoshop and Illustrator software (Adobe).

### Hydrogen-Deuterium Exchange Mass Spectrometry (HDX-MS)

Protein samples were diluted to 58.9:118 μM GINIP:Gαi3-GTPγS or 58.9 μM GINIP alone in HDX-MS buffer (20 mM HEPES pH 7.4, 150 mM NaCl, 10 mM MgCl_2_, 1 mM DTT) and kept in a temperature controlled drawer at 4 °C of a LEAP HDX robotics set-up for the duration of the sampling scheme (approximately 24 h). All protein handling steps from this point on were automated, including a staggered sampling scheme to optimize LC-MS runtime. The entire process of exchange, digestion, separation and MS detection for all samples was carried out in ∼24 h. For each condition tested, 3.8 μl of protein were taken from the stock vials and added to empty vials kept in a temperature controlled drawer at 25 °C. 52.2 μl of room temperature D_2_O-based HDX-MS buffer (or H_2_O-based buffer for non-exchanged controls) were dispensed to protein samples and incubated for 10 s, 100 s, 1000 s, or 10000 s. Exchange reactions were stopped by mixing 50 μl of the samples with 50 μl of pre-chilled quench buffer (100 mM potassium phosphate, pH 2.66) in a 4 °C drawer. 95 μl of the quenched reactions were immediately injected for in-line digestion in a pepsin column (Waters, 186007233) followed by LC-MS analysis of peptides (ACQUITY UPLC M-Class – Synapt G2Si HD MS, Waters). Exchange reactions for each protein sample were performed in at least triplicate in a single experiment. Peptide identification was performed using ProteinLynx Global Server (PLGS) v 3.0.1 software (Waters). Undeuterated samples were searched against a database with the sequences of human GINIP and rat Gαi3. Identified peptides were imported to DynamX v 3.0 software for analysis of deuterium uptake. Peptides were manually curated to ensure they matched parameters of retention time and ion mobility for the peak of the isotopic distribution across different exposures and replicate samples for each charge state of identified peptide. Deuterium uptake was determined by subtracting the centroid mass of undeuterated peptides from those of deuterated peptides. Relative fractional uptake was calculated by dividing the average increase in mass (Daltons) of deuterated peptides by the total number of backbone hydrogens available for exchange. Heat maps were generated representing per-residue relative fractional uptake for GINIP:Gαi3 or GINIP alone as well as the difference between the two states to identify regions of protection or de-protection.

### Differential Scanning Fluorimetry (DSF) thermal shift assay

Thermal shift assays were carried out in 96-well semi-skirted PCR plates using purified, C-terminally His-tagged GINIP. GINIP proteins were diluted on ice to 8.3 μM in PBS and 10 μl of this diluted sample were added to the plate at room temperature. 5000X SYPRO Orange protein stain (Life Technologies, S6650) was diluted 100 fold in PBS and 15 μl of this diluted reagent were added to the wells containing protein samples at room temperature (final volume of 25 μl; 5 μM recombinant GINIP, 20X SYPRO Orange) and spun down for one minute at 200 x g. Fluorescence data for SYPRO Orange signal was collected with a ViiA 7 Real-Time PCR System (Applied Biosystems) and QuantStudio Real-Time PCR Software v1.2 using 470 ± 15 nm excitation and 586 ± 10 nm emission filter. The plates were held at 25 °C for 2 minutes to stabilize the sample temperature, after which initial fluorescence was measured. Subsequent reads were taken at each temperature interval (0.5 °C steps) after 20 seconds of temperature stabilization for each step. Measurements were performed in technical triplicates and averaged for each independent experiment. Melting curves were generated by reads spanning 25-55 °C and are normalized to the maximum and minimum intensities measured for each protein condition. The thermal denaturation temperature (T_m_) was determined by plotting normalized intensity as a function of temperature and fitting the transition region to a Boltzman sigmoid. The T_m_ is the midpoint between the baseline intensity (*I_b_*, nondenaturated) and the peak intensity plateau value, (*I_p_*, maximum denaturation) following the equation below.

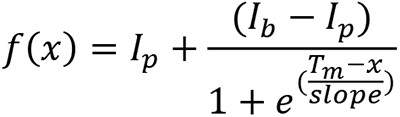

### Measurement of the associations of GINIP and Gαi in HEK293T cells by BRET

Approximately 400,000 HEK293T cells were seeded on each well of 6-well plates coated with 0.1% (w/v) gelatin, and transfected ∼24 hr later using the calcium phosphate method with plasmids encoding the following constructs as indicated in the figures (DNA amounts in parentheses): GABA_B_R1a (0.2 μg), GABA_B_R2 (0.2 μg), Gαi3-Nluc(a/b) (0.05 μg), Gαi3-YFP(a/b) (0.5 μg), GINIP-YFP WT (0.5 μg), GINIP-YFP W139A (0.5 μg), GINIP-Nluc WT (0.05 μg), and GINIP-Nluc W139A (0.05 μg). Total DNA amount per well was equalized by supplementing with empty pcDNA3.1 as needed. Cell medium was changed 6 h after transfection, and approximately 16-24 h after transfection, cells were washed and gently scraped in room temperature PBS, centrifuged (5 min at 550 × *g*), and resuspended in BRET buffer (140 mM NaCl, 5 mM KCl, 1 mM MgCl_2_, 1 mM CaCl_2_, 0.37 mM NaH_2_PO_4_, 24 mM NaHCO_3_, 10 mM HEPES and 0.1% glucose, pH 7.4) at a concentration of ∼10^6^ cells/ml. ∼25,000 cells/well were added to a white opaque 96-well plate (Opti-Plate, PerkinElmer Life Sciences, 6005290) and mixed with the nanoluciferase substrate Nano-Glo (Promega, N1120, final dilution 1:200) before measuring luminescence. Luminescence signals at 450 ± 40 and 535 ± 15 nm were measured at 28 °C every 0.96 s in a BMG Labtech POLARStar Omega plate reader and BRET was calculated as the ratio between the emission intensity at 535 nm divided by the emission intensity at 450 nm. Kinetic traces are represented as an increase in BRET after subtraction of the baseline signal measured for 30 s before GPCR stimulation (ΔBRET).

### Protein structure modelling and visualization

The folding model of GINIP (Uniprot #Q9UPV7) was extracted from AlphaFold 2.0 (Jumper et al., 2021). The folding models of the GINIP:Gαi complex and GRIN3:Gαi complex were generated using ColabFold (Jumper et al., 2021; Mirdita et al., 2022) accessed using UCSF ChimeraX 1.4 (Pettersen et al., 2004). ColabFold is an open-source, user-accessible program using a Google Colaboratory notebook that leverages features of AlphaFold 2.0 with improvements on specific steps. Full-length sequences for human GINIP (Uniprot #Q9UPV7) and human Gαi1 (Uniprot #P63096) were input into the notebook and the ensemble, best-fit model based on the best five output models based on pLDDT (predicted local difference test) was used for presentation. pLDDT is a per-residue confidence metric measured on a scale of 0-100, pLDDT values were overlaid on structures shown in **Fig S5A** (Jumper et al., 2021). For images of G proteins bound to effectors and effector-like molecules shown in **Fig. 4B**, the following PDB IDs were used: KB-1753:Gαi1 (PDB ID: 2G83), GN13:Gαs (PDB ID: 7BPH), PDEγ:Gαt/i (PDB ID: 1FQJ), and PLCβ:Gαq (PDB ID: 7SQ2) (Dai et al., 2022; Johnston et al., 2006; Slep et al., 2001; Waldo et al., 2010). Images of structures were captured using Open-Source PyMOL 1.2r1.

### Measurement of GINIP intramolecular BRET upon G protein binding in cell lysates

Purified Gαi3 loaded with GDP or GTPγS was mixed to lysates of cells expressing GINIP dually fused to a BRET donor (Nluc) and a BRET acceptor (YFP) to detect changes in the distance or relative orientation of the BRET pair. Gαi3 was loaded with GDP or GTPγS by incubating the protein at 30 °C with 150 μM GDP or GTPγS in G protein storage buffer (20 mM Tris-HCl, 20 mM NaCl, 1 mM MgCl_2_, 1 mM DTT, 5 % glycerol (v:v)) for 3 hours. Nucleotide-loaded G proteins were aliquoted and stored at −80 °C. G protein stocks were serially diluted in G protein storage buffer supplemented with 150 μM of GDP or GTPγS to obtain the following concentrations: 0. 1.25, 2.5, 5, and 10 μM. Ten μl of these dilutions or 10 μl of G protein storage buffer without nucleotides were pipetted into the corner of a white opaque 96-well plate (Opti-Plate, PerkinElmer Life Sciences, 6005290). Ten 10 μl of 0.1 mM Coelenterezine 400a diluted in BRET buffer (140 mM NaCl, 5 mM KCl, 1 mM MgCl_2_, 1 mM CaCl_2_, 0.37 mM NaH_2_PO_4_, 24 mM NaHCO_3_, 10 mM HEPES and 0.1% glucose, pH 7.4) were pipetted into the opposite corner of the well without mixing with the G protein. Then, 80 μl of cell lysates expressing different GINIP constructs, prepared as explained next, were added to the wells. Briefly, approximately 400,000 HEK293T cells were seeded on each well of 6-well plates coated with 0.1% (w/v) gelatin, and transfected ∼24 hr later with 50 ng of plasmids encoding the following constructs: p3xFLAG-CMV-14-Nluc-GINIP-YFP, p3xFLAG-CMV-14-Nluc(31)-GINIP-YFP, p3xFLAG-CMV-14-YFP-GINIP-Nluc, and p3xFLAG-CMV-14-YFP(31)-GINIP-Nluc. Cells were lysed by addition of Triton X-100 to a final concentration of 0.1% (v/v) and incubated at room temperature for 4 minutes. After addition to the lysates to the wells of the white opaque 96-well plate, luminescence was measured in a BMG Labtech POLARStar Omega plate reader. Luminescence signals at 450 ± 40 and 535 ± 15 nm were measured three times at 28 °C at 30 s increments in a BMG Labtech POLARStar Omega plate reader. BRET was calculated as the ratio between the emission intensity at 535 nm divided by the emission intensity at 460 nm for the three individual measurements and then averaged. The value for the BRET ratio of G protein storage buffer without nucleotides was subtracted from all values. ΔBRET was calculated as the difference between the BRET ratio of samples including G protein subtracted from the value of the BRET ratio calculated for G protein storage buffer with nucleotide alone.

### cAMP measurements in HEK239T cells by BRET

Approximately 400,000 HEK293T cells were seeded on each well of 6-well plates coated with 0.1% (w/v) gelatin, and transfected ∼24 hr later using the calcium phosphate method with plasmids encoding the following constructs as indicated in the figures (DNA amounts in parentheses): Nluc-EPAC-VV (0.05 μg), GABA_B_R1a (0.2 μg), GABA_B_R2 (0.2 μg), and GINIP-FLAG (2 μg). Total DNA amount per well was equalized by supplementing with empty pcDNA3.1 as needed. Cell medium was changed 6 h after transfection, and approximately 16-24 h after transfection, cells were washed and gently scraped in room temperature PBS, centrifuged (5 min at 550 × *g*), and resuspended in BRET buffer (140 mM NaCl, 5 mM KCl, 1 mM MgCl_2_, 1 mM CaCl_2_, 0.37 mM NaH_2_PO_4_, 24 mM NaHCO_3_, 10 mM HEPES and 0.1% glucose, pH 7.4) at a concentration of ∼10^6^ cells/ml. Approximately 25,000 cells/well were added to a white opaque 96-well plate (Opti-Plate, PerkinElmer Life Sciences, 6005290) and mixed with the nanoluciferase substrate Nano-Glo (Promega, N1120, final dilution 1:200) before measuring luminescence. Luminescence signals at 450 ± 40 and 535 ± 15 nm were measured at 28 °C every 4 s in a BMG Labtech POLARStar Omega plate reader and BRET was calculated as the ratio between the emission intensity at 535 nm divided by the emission intensity at 450 nm. Since the Nluc-EPAC-VV construct reports cAMP binding as a decrease in BRET, results were processed as the inverse of the BRET ratio (BRET^-1^) to make it more intuitive. After subtraction of a basal signal measured for 30 s before stimulation with forskolin (ΔBRET^−1^), results were normalized to the maximum response of forskolin detected prior to the addition of GPCR agonists (cAMP (normalized ΔBRET^−1^). For calculation of the agonist-mediated inhibition of cAMP induced by forskolin “inhibition (%FSK),” the average of the last six time points of ΔBRET^−1^ curves were used. Baseline ΔBRET^−1^ values from cells not treated with forskolin were subtracted from values obtained from forskolin and agonist treated or forskolin treated conditions. Inhibition (%FSK) was represented as the ratio of forskolin and agonist treated divided by forskolin treated.

### Free Gβγ measurements in HEK293T cells by BRET

Approximately 400,000 HEK293T cells were seeded on each well of 6-well plates coated with 0.1% (w/v) gelatin, and transfected ∼24 hr later using the calcium phosphate method with plasmids encoding the following constructs as indicated in the figures (DNA amounts in parentheses): VN-Gγ2 (0.1 μg), VC-Gβ1 (0.1 μg), masGRK3ct-Nluc (0.1 μg), Gαi3 WT (0.5 μg), GABA_B_R1a (0.2 μg), GABA_B_R2 (0.2 μg), RGS8 (0.5 μg), RGS12 (0.5 μg), and GINIP-FLAG (2 μg). Luminescence measurements were carried out as in “*cAMP measurements in HEK239T cells by BRET*,” except signals were recorded every 0.24 s. Results were presented as increase in BRET after subtraction of the basal signal measured for 30 s before any stimulation (ΔBRET (baseline)). For the calculation of response amplitudes, the difference between the raw BRET ratio before and 60 s after agonist stimulation was calculated. For the presentation and calculation of G protein deactivation rates, the minimum BRET value reached after the addition of antagonist (plateau signal) was subtracted from the raw BRET ratio of each timepoint (recovery corrected ΔBRET), and each resulting value was scaled to as the percentage of the maximal BRET value right before the addition of antagonist (“% Maximum response”). G protein deactivation rate constants (*k*) were determined by fitting the recovery-corrected ΔBRET values after the addition of antagonist to a one-phase decay equation (*Y* = (*Y*_0_ − *Plateau* × *e*^(−*kX*)^ + *Plateau*) in Prism 9 (Graphpad), where *Y_0_* is the starting value right before the addition of antagonist (constrained to 100) and *Plateau* is the near-zero minimum estimated by the fit.

### Statistical analyses

Unless otherwise indicated, experiments were independently repeated a minimum of three times. For experiments displaying pooled data, individual data points and/or mean ± S.E.M or ± SD is depicted with the exception of curves in **Fig S4** where only the mean is represented. For other experiments, like immunoblot images, one representative result is presented. All statistical comparisons were calculated in GraphPad Prism 9. A Student’s t-test was used in cases where two conditions were compared. One-way ANOVA with correction for multiple comparisons using Tukey post-hoc analysis was used in cases where 3 or more conditions were compared. In the case where differences in quantitative variables for two independent variables were compared, as in **Fig. 2C**, a two-way ANOVA was used without correction for multiple comparisons using Fisher’s LSD test.

## ACKNOWLEDGEMENTS

This work was primarily supported by NIH grant R01NS117101 (to M.G-M.). A.L. is supported by a F31 Ruth L. Kirschstein NRSA Predocotral Fellowship (F31NS115318). We thank the following investigators for providing DNA plasmids: N. Lambert (Augusta University, Augusta, GA), K. Martemyanov (UF Scripps, Jupiter, FL), C. Dessauer (University of Texas Health Science Center at Houston, TX), M. Linder (Cornell University), P. Slessinger (Mount Sinai NY).

## AUTHOR CONTRIBUTIONS

A.L. and M.Z. conducted experiments. A.L. and M.G-M. designed experiments and analyzed data. S.J.E. provided training and technical support for conducting and analyzing mass spectrometry experiments. A.L. and M.G-M. wrote the manuscript with input from all authors. M.G-M. conceived and supervised the project.

---

**Figure S1.**
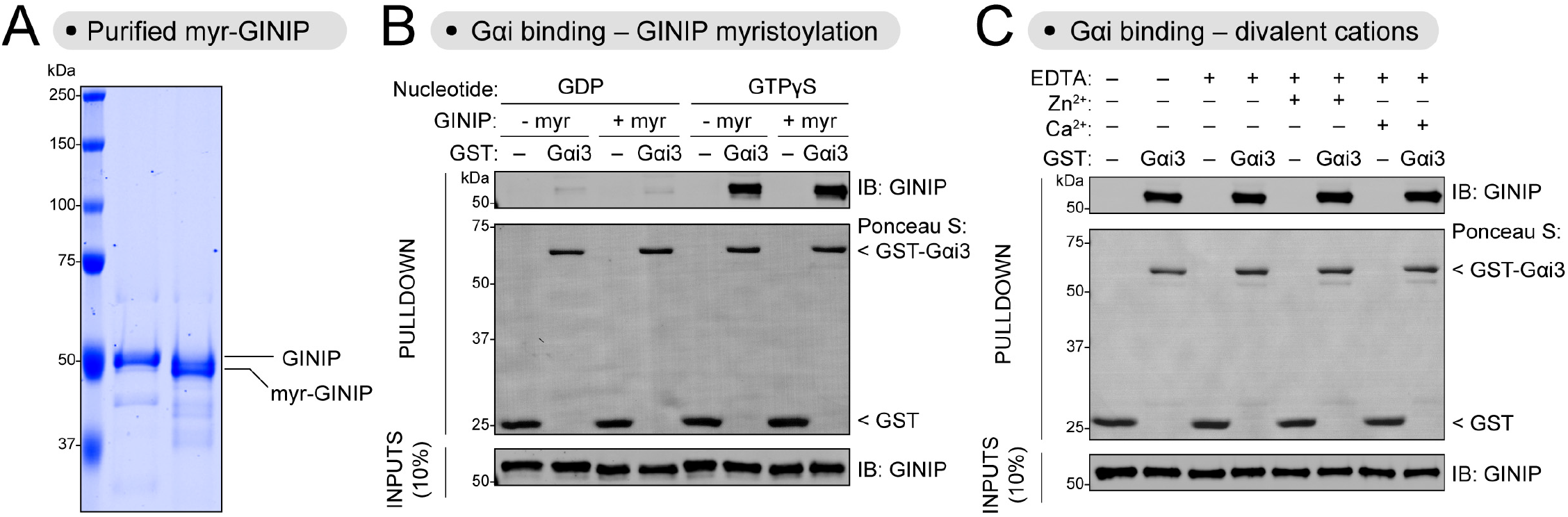
Neither GINIP myristoylation nor presence of divalent cations affects binding to Gαi3. **(A)** His-tagged non-myristoylated and myristoylated-GINIP (myr-GINIP) purified from bacteria were run in an 8% SDS-PAGE gel supplemented with 4M urea before Coomassie staining. Two μg of each protein were loaded. Myr-GINIP displays a downward shift compared to GINIP. **(B)** Myristoylation of GINIP does not affect binding to Gαi3. Purified GINIP or myr-GINIP were incubated with GST or GST-Gαi3 immobilized on glutathione-agarose beads in the presence of GDP or GTPγS as indicated. Bead-bound proteins were detected by Ponceau S staining or by immunoblotting (IB) as indicted. **(C)** Divalent cations do not affect the binding of GINIP to Gαi3. Purified GINIP was incubated with GST or GST-Gαi3 immobilized on glutathione-agarose beads in buffers with the indicated additives (Zn^2+^ = ZnCl_2_, Ca^2+^ = CaCl_2_). All conditions contained GTPγS. Bead-bound proteins were detected by Ponceau S staining or by immunoblotting (IB). Results are representative of n ³ 3 experiments.

**Figure S2.**
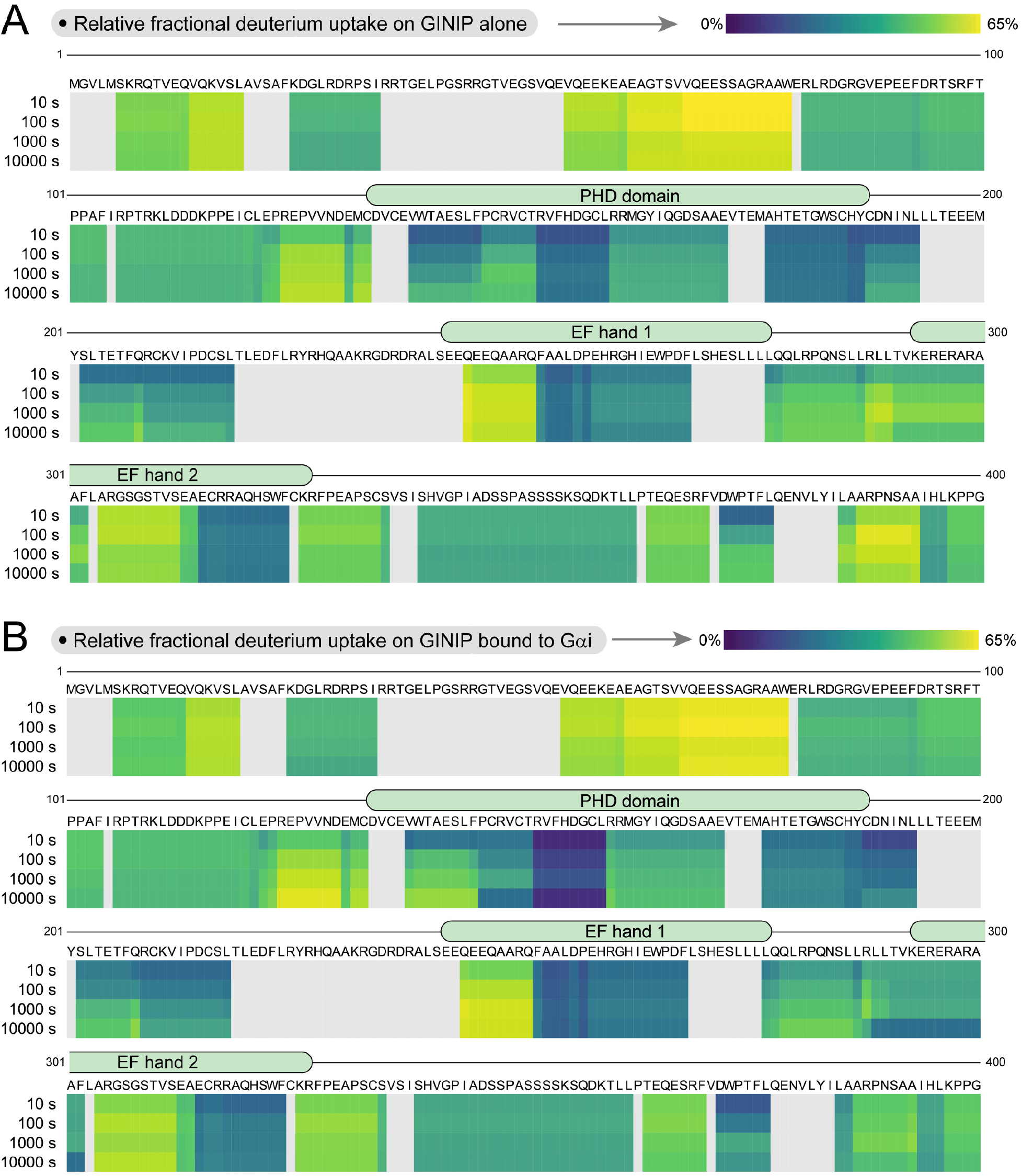
Relative fractional uptake of GINIP alone or in complex with Gαi3. **(A, B)** Stacked heat maps of relative fractional uptake of deuterium by GINIP peptides from GINIP alone (A) or the GINIP:Gαi complex (B) at different times (10 s, 100 s, 1000 s, 10000 s), where navy blue is no exchange and yellow is maximal exchange measured. Grey indicates regions without peptide coverage. Data are the mean mean relative fractional uptake an experiment run in quadruplicates.

**Figure S3.**
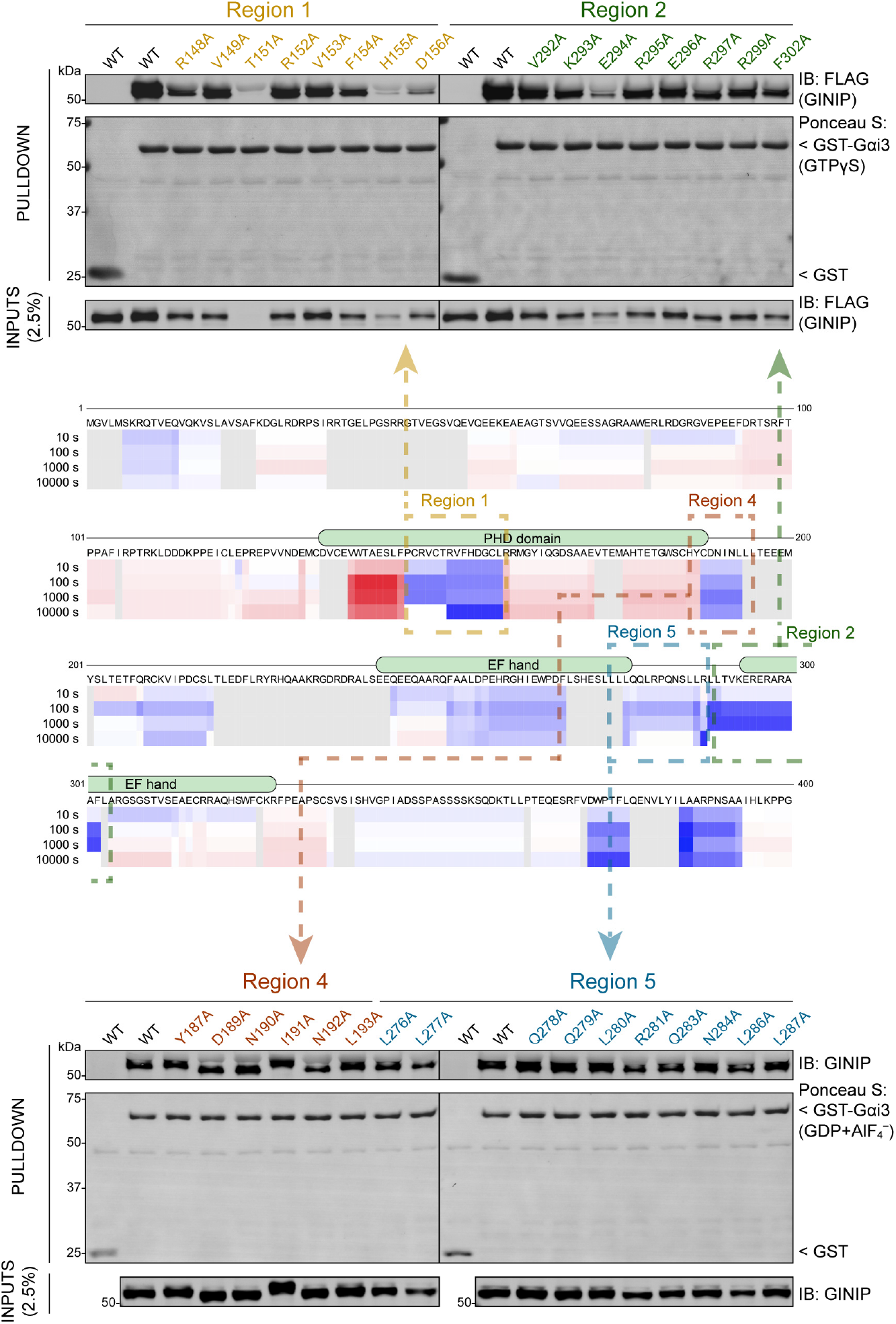
Mutation of residues in several regions of GINIP with decreased deuterium uptake does not disrupt binding to active Gαi3. *Center,* Stacked heat map of the difference in relative fractional uptake of deuterium by GINIP peptides from the GINIP:Gαi complex relative to GINIP alone at different times (10 s, 100 s, 1000 s, 10000 s), where blue is a decrease in deuterium uptake and red is an increase. White means no difference and grey indicates regions without peptide coverage. Boxes indicate the regions with the largest (Regions 1 and 2) or moderate decreases in deuterium uptake (Region 4 and 5). This heat map is reproduced from Fig 2. *Top & Bottom,* mutation of indicated single residues in Regions 1, 2, 4, and 5 did not reduce binding for those mutants that were expressed. Lysates of HEK293T cells expressing the indicated GINIP mutants were incubated with GST or GST-Gαi3 immobilized on glutathione-agarose beads in the presence of GDP, GDPꞏAlF_4_^-^, or GTPγS, as indicated. Bead-bound proteins were detected by Ponceau S or by immunoblotting (IB). All protein electrophoresis results are representative of n ³ 3 experiments.

**Figure S4.**
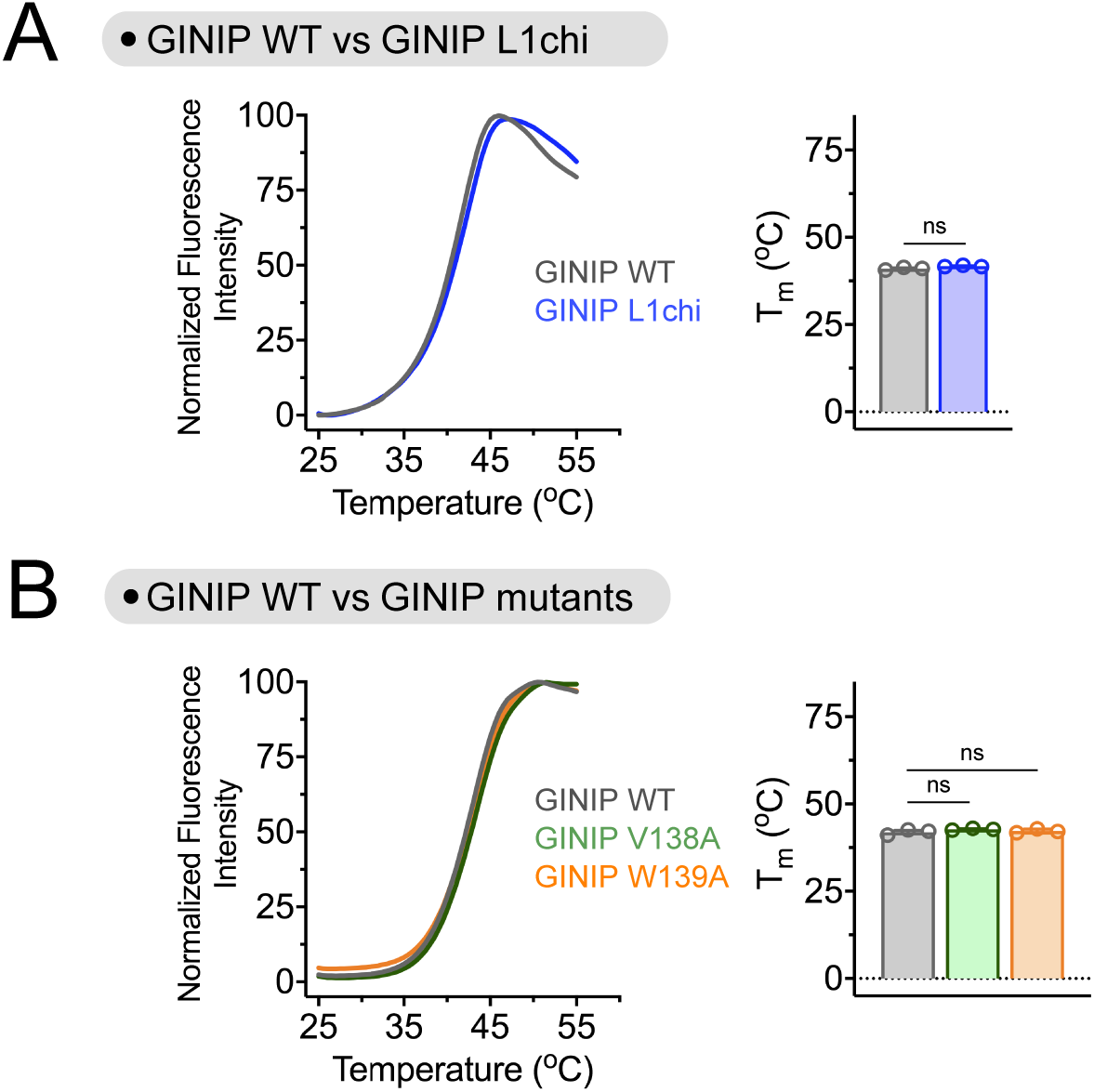
GINIP mutant proteins display the same thermal stability as GINIP WT. **(A,B)** GINIP L1chi, GINIP V138A, and GINIP W139A have thermal stability similar to GINIP WT as measured by Differential Scanning Fluorimetry (DSF). *Left graphs,* Thermal denaturation curves comparing GINIP L1chi (blue, A) or GINIP V138A (green, B) and W139A (orange, B) to GINIP WT (grey, A and B). SYPRO Orange Fluorescence was measured to track denaturation as temperature of protein samples was increased over time. Mean, n=3. *Right graphs,* Quantified melting temperature (T_m_) of GINIP mutants compared to GINIP WT based on the sigmoid of the denaturation curves. Mean ± S.E.M., n=3, ns = not significant.

**Figure S5.**
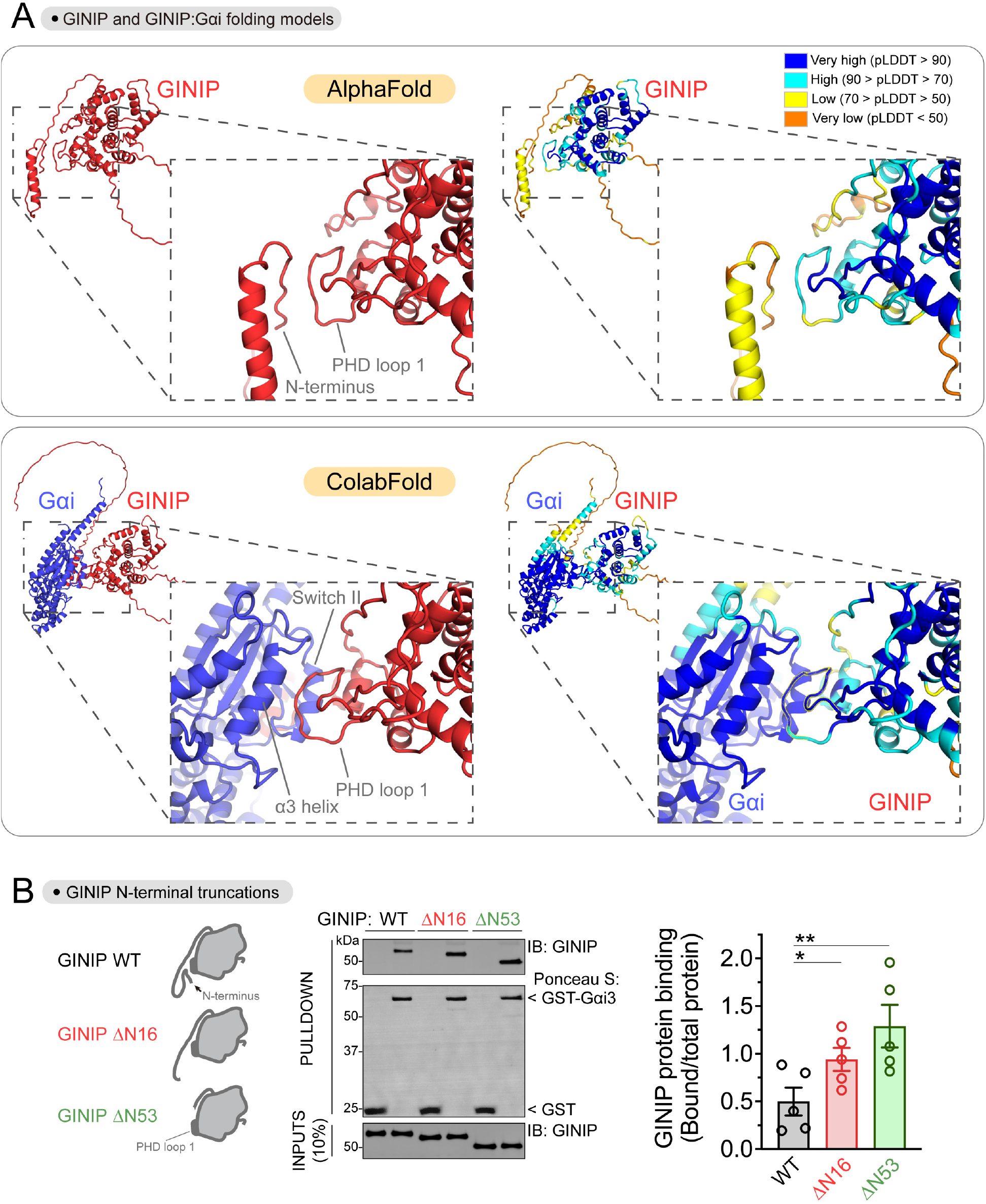
Putative intramolecular contact between the N-terminus of GINIP and the Loop 1 of the PHD domain. **(A)** The N-terminus of GINIP might be in the vicinity of the Loop 1 of the PHD domain in the absence of Gαi but not when bound to the G protein. The AlphaFold 2.0 model of GINIP alone is shown on the top, and the ColabFold model of GINIP in complex with Gαi is shown on the bottom. GINIP and Gαi are colored red and blue, respectively on the left, whereas the panels on the right are colored based on predicted local difference test (pLDDT) scores, a per-residue confidence metric measured on a scale of 0-100. **(B)** N-terminally truncated GINIP constructs display modest increases in binding to Gαi3. *Left,* schematic representation of N-terminal truncations of GINIP removing residues 1-16 (ΔN16) or 1-53 (ΔN53) and the resulting exposure of the PHD Loop 1 G protein binding site. *Center,* Purified His-tagged GINIP WT, GINIP ΔN16, or GINIP ΔN53 were incubated with GST or GST-Gαi3 immobilized on glutathione-agarose beads in the presence of GTPγS. Bead-bound proteins were detected by Ponceau S staining or by immunoblotting (IB). A representative experiment of n=5 is shown on the left, and a graph with the quantification of binding is shown on the right. Mean ± S.E.M., *p<0.05, **p<0. 01, paired one-way ANOVA corrected for multiple comparisons (Tukey).

**Figure S6.**
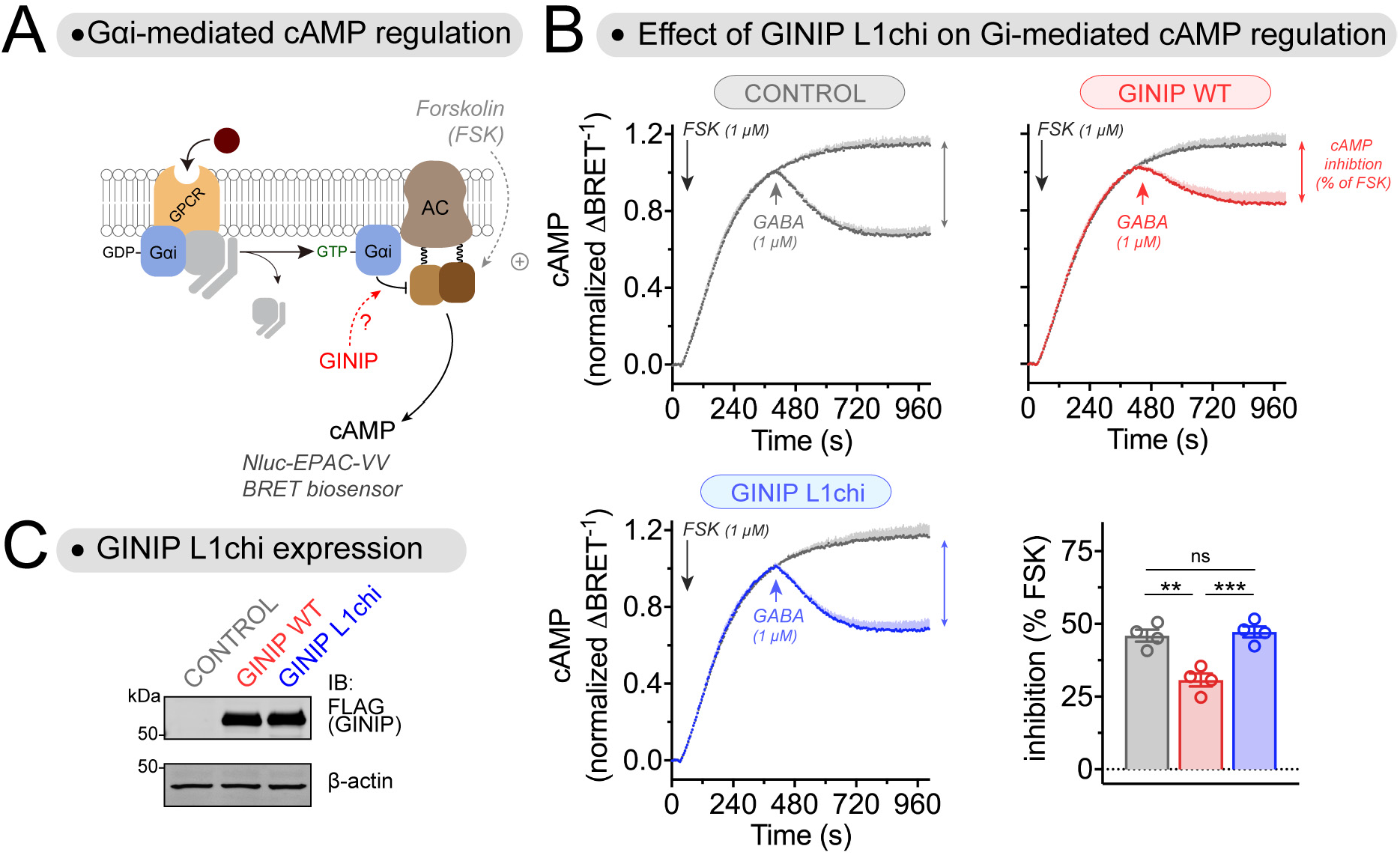
GINIP L1chi fails to regulate cAMP cellular levels upon GPCR stimulation. **(A)** Diagram of GPCR-mediated activation of Gαi-GTP and subsequent regulation of cAMP levels in cells monitored by BRET. **(B)** Replacement of the Loop 1 of the PHD domain in the GINIP L1chi construct prevents the blockade of cAMP inhibition upon stimulation of GABA_B_R observed with GINIP WT. Kinetic BRET measurements of cAMP levels were carried out in HEK293T cells expressing the GABA_B_R without GINIP (gray) or expressing GINIP WT (red) or GINIP L1chi (blue). Cells were treated with forskolin (FSK) and GABA as indicated. Quantification of the inhibition of FSK-stimulated cAMP upon stimulation of GABA_B_R with GABA is shown in the bar graph on the bottom left corner. Mean ± S.E.M., n=4. ns = not significant, **p<0.01, ***p<0.001, one-way ANOVA corrected for multiple comparisons (Tukey). **(C)** Representative immunoblotting (IB) result confirming equal expression of GINIP WT and GINIP L1chi in the cells used for the experiments shown in (B).

**Figure S7.**
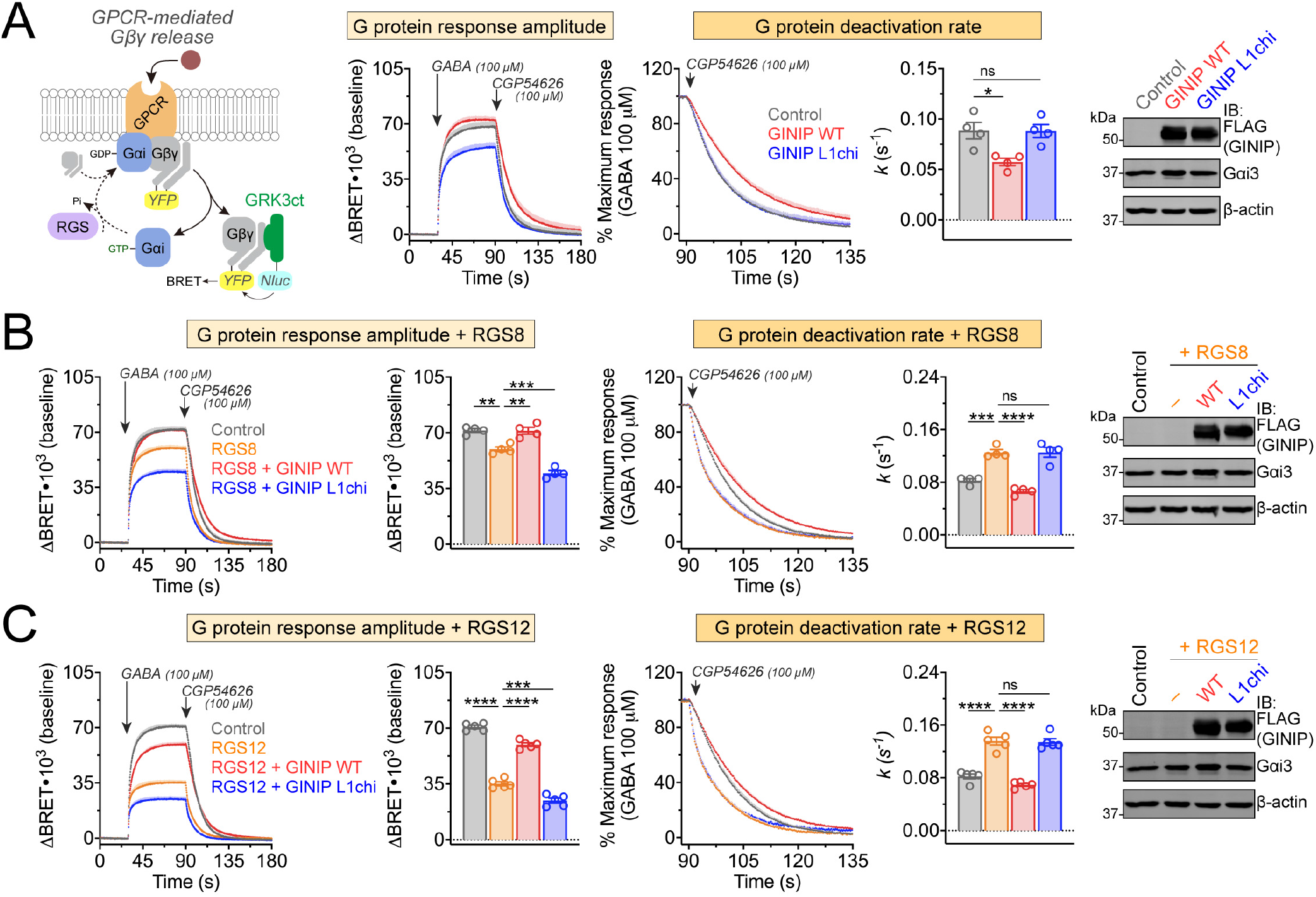
GINIPL1chi fails to regulate Gβγ responses in cells upon GPCR stimulation. **(A)** Expression of GINIP Loop 1 chimera (L1chi) prevents the enhancement of Gβγ signaling upon stimulation of GABA_B_R observed with GINIP WT. *Left,* diagram of G protein activation/deactivation cycle and BRET-based detection of free Gβγ. *Center,* kinetic BRET measurements were carried out in HEK293T cells expressing the GABA_B_R without GINIP (grey), or expressing GINIP WT (red) or GINIPL1chi (blue). Cells were treated with GABA and CGP54626 as indicated. *Right,* G protein deactivation rates were determined by normalizing the BRET data to maximum response and fitting the post-antagonist data to an exponential decay curve to extract rate constant values (*k*). Mean ± S.E.M., n=4. ns = not significant, *p<0.05, one-way ANOVA corrected for multiple comparisons (Tukey). A representative immunoblotting (IB) result confirming equal expression of GINIP WT, GINIPL1chi, and Gαi3 in these experiments is shown on the right. **(B)** Replacement of Loop 1 of GINIP ablates RGS8-mediated regulation of Gβγ signaling upon stimulation of GABA_B_R observed with GINIP WT. BRET experiments were carried out and analyzed as in (A) with cells expressing RGS8 alone (orange), RGS plus GINIP WT (red), RGS plus GINIPL1chi (blue), or neither RGS8 nor GINIP (grey). Quantification of G protein response amplitude was determined 1 minute after agonist stimulation. Mean ± S.E.M., n=4. ns = not significant, **p<0.01, ***p<0.001, ****p<0.0001, one-way ANOVA corrected for multiple comparisons (Tukey). **(C)** Replacement of Loop 1 of GINIP ablates RGS12-mediated regulation of Gβγ signaling upon stimulation of GABA_B_R observed with GINIP WT. BRET experiments were carried out and analyzed as in (A) with cells expressing RGS12 alone (orange), RGS plus GINIP WT (red), RGS plus GINIPL1chi (blue), or neither RGS8 nor GINIP (grey). Quantification of G protein response amplitude was determined 1 minute after agonist stimulation. Mean ± S.E.M., n=5. ns = not significant, ***p<0.001, ****p<0.0001, one-way ANOVA corrected for multiple comparisons (Tukey). A representative immunoblotting (IB) result confirming equal expression of GINIP WT, GINIP mutants, and Gαi3 in these experiments is shown on the right.

**Figure S8.**
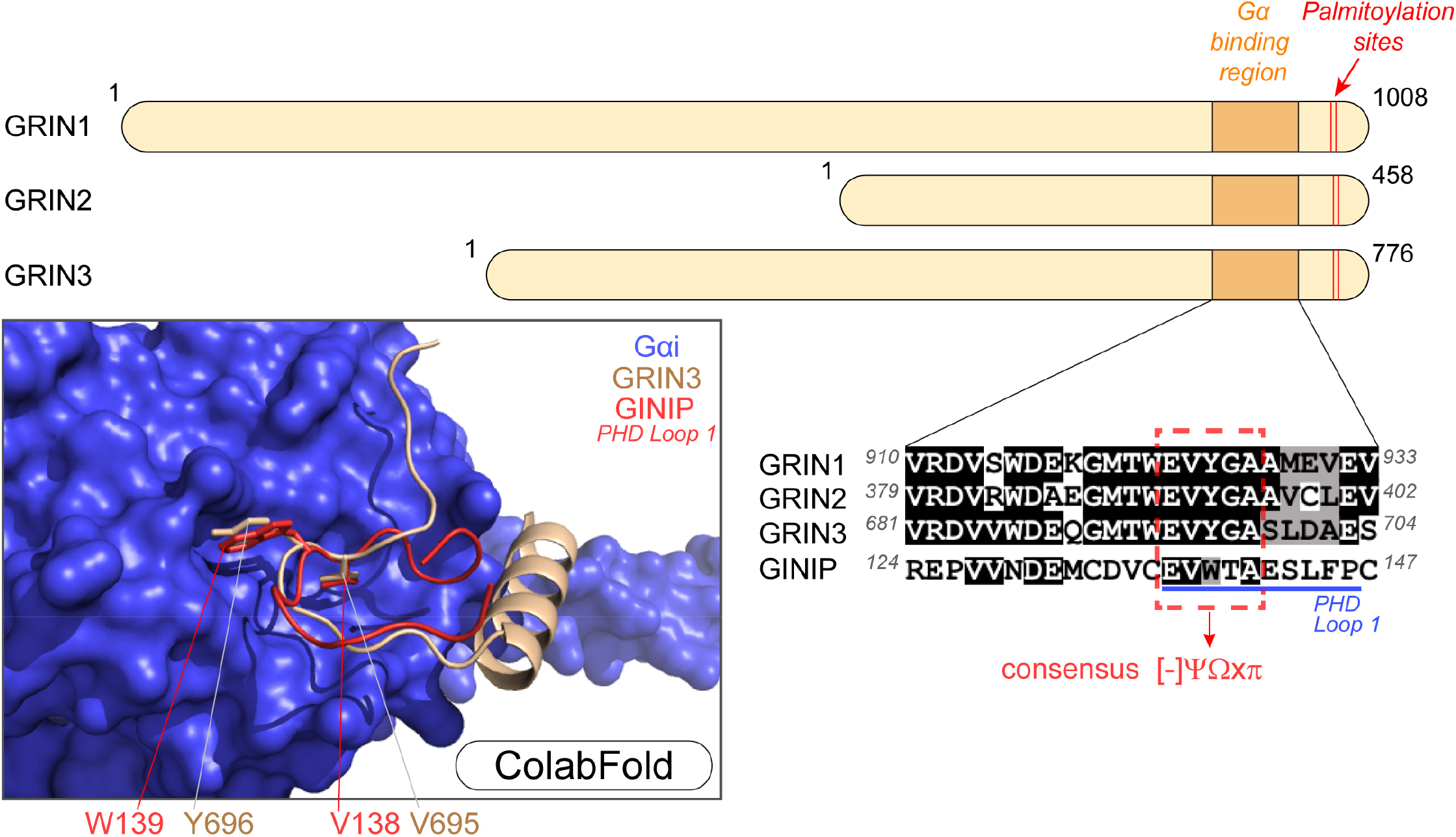
Predicted similarity between Gα binding modes of GINIP and GRIN. *Top and bottom right,* sequence alignment of GINIP’s PHD loop 1 and a G protein binging region of GRIN family proteins (GRIN1, GRIN2 and GRIN3) suggests a consensus motif for Gαi binding. Sequence alignment suggests a potential short consensus motif [–]-Y-W-x-p, where [–] is a negatively charged residue, Y is a residue with an aliphatic chain, W is an aromatic residue, x is any residue, and p is a residue with a small side-chain. *Bottom left*, image of protein folding model generated in ColabFold for GINIP (red) bound to Gαi (blue) overlaid with an independently generated model of GRIN3 (tan) bound to Gαi. Amino acids V695 and Y696 located in the consensus motif of GRIN proteins shared with GINIP dock into Gαi with a pose similar to that observed for the equivalent amino acids of GINIP, V138 and W139, which are required for G protein binding and regulation.

